# Uncertainty-Aware Gene Rankings Reveal Key Players in Coexpression Networks

**DOI:** 10.64898/2025.12.13.694156

**Authors:** Sugyani Mahapatra, Nilesh Anantha Subramanian, Manikandan Narayanan

## Abstract

**Motivation:** Key genes of a biological system are often prioritized by computing network science measures on a coexpression network inferred from transcriptomic data. But human population heterogeneity and modest sample sizes introduce uncertainty in the inferred coexpression network. Earlier studies have estimated this uncertainty using bootstrap resampling or similar approaches, but fewer have investigated how it propagates to downstream network analyses and affects gene prioritization.

**Methods and Results:** We present a systematic workflow to propagate network uncertainty to downstream measures such as degree/PageRank centrality, with the goal of producing robust gene scorings/rankings. We specifically propose uncertainty-aware scorings, `BooNS` and `BPNS`, which utilize the spread of a centrality measure across bootstrapped coexpression networks to prioritize stable central genes. Across several (semi-)simulated and real-world (GTEx) datasets, `BooNS` and `BPNS` recover reference or tissue-specific genes significantly better than other traditional centrality rankings. This performance gap highlights the long-overdue adoption of uncertainty-aware gene ranking for stable biological inference.

**Availability and Implementation:** Code and supplemental data are available at https://github.com/BIRDSgroup/Bootstrap-based_Node_Scorings_BNS, and https://tinyurl.com/BNS-suppl-data respectively.

## 1. Introduction

Gene-gene coexpression networks inferred from transcriptomics data aim to capture co-regulatory and functional relationships among various genes in a complex biological system; and analyzing them using network science measures such as node centrality, “hub”ness, clustering coefficient, and k-core number have been successful in revealing key regulatory genes and new insights into structural/functional organization of the system (2; 36; 29). But, relatively modest sample sizes of many transcriptomic studies, along with heterogeneity in the human population (i.e., individual-to-individual variation of extraneous factors that influence a gene expression trait), can lead to uncertainty about the “true” gene coexpression network.

Bootstrap resampling or similar analysis have been used to address this uncertainty in network inference from genomic data. For instance, several studies have repeatedly inferred a network on different resampled subsets of the observed data, and aggregated information across these networks to estimate the uncertainty and conversely the confidence level of an inferred network edge (e.g., (18; 26; 9; 6)). Certain studies have also propagated uncertainty in the inferred network (edges) to downstream network measures such as node degree in order to choose a network inference algorithm resulting in stable degrees (15), centrality indices to enable pairwise comparison of node centralities in psychological networks (12), edge/node membership in modules to assess stability of a clustering (33), densities of certain subgraphs to identify robust network motifs (5), and degree distribution to estimate mean or outlier degree across all nodes (19). But these methods have not addressed how uncertainty impacts the ranking or prioritization of all genes in a biological network based on per-gene importance measures such as centrality – this ranking is a critical step towards prioritizing experimental and functional validation of different “hub” genes.

We take a step forward in this direction by presenting a systematic workflow that estimates and propagates uncertainty in the inferred coexpression network to downstream measures such as degree and PageRank centrality, with the main goal of producing a robust (uncertainty-aware) gene ranking. Given the distribution of a network science measure across multiple bootstrapped coexpression networks with mean *µ* and variance *σ*^2^, we propose different uncertainty-aware gene rankings, namely, “*ESTimator-based Node Scores* (EstNS)”, “*BOotstrap Offset-based Node Scores* (`BooNS`)” of the form *µ − cσ* (for different values of *c*) and “*Bootstrap Percentile-based Node Scores* (`BPNS`)” of the form 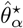 (representing the *α*^th^ percentile) to obtain robust rankings of genes. Genes with high values of *µ* and low *σ* indicate high stability in their centrality values – they not only constitute core hub genes in the system, but also show less sensitivity to noise in the network revealed by the random bootstrap resampling process (39). Our proposed rankings score such genes higher than the other genes either by penalizing the variability in their centrality measures as in EstNS and `BooNS`, or with more conservative rankings using bootstrap percentiles as in `BPNS`.

The central contribution of our study is not only to propose simple penalized scores that incorporate bootstrap-based uncertainty into downstream network analysis, but also to carefully evaluate their efficacy in yielding a robust global ranking of genes. Specifically, our work makes these contributions.

1. We present a framework that estimates network uncertainty using bootstrap and more importantly propagates it to downstream network science measures such as degree and PageRank centrality using uncertainty-aware scorings such as bootstrap offset-based node scores (`BooNS`) of the form *µ −cσ* and bootstrap percentile confidence interval based node scores (`BPNS`) of the form 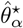 .
2. We systematically assess the performance of gene rankings based on these proposed scorings with respect to reproducibility, replicability, and tissue specificity across simulated, semi-simulated and real-world (multi-tissue GTEx) bulk transcriptomic datasets, and show it mostly outperforms the baseline rankings that do not incorporate uncertainty.
3. Furthermore, we analyze fluctuations in our gene scores and rankings for different dataset sizes to provide sample size recommendations.

Given the promising results we observe from these comprehensive datasets and analyses, our systematic approach for quantifying and propagating network uncertainty can be applied to study other measures (e.g., clustering coefficients or connectivity measures based on paths or components) and other transcriptomic datasets (e.g., single-cell or spatial transcriptomic datasets) as well.

## 2. Methods

### 2.1. Our Proposed Scorings

We propose the use of variance in estimates of a network measure across multiple bootstrapped coexpression networks for propagation of uncertainty. Repeated application of the bootstrapping process on the observed gene expression dataset generates several (*B*) “*bootstrap resampled datasets*” (sample-size-matched datasets obtained via sampling with replacement from the observed dataset), and a coexpression network is then inferred from each resampled dataset (see Fig. 1A). For a gene *g*, its centrality is computed in each of these *B* networks, and the mean and standard deviation of the resulting *B* values is denoted by *µ*[*g*] and *σ*[*g*] respectively (with *µ* and *σ* referring to the vector of such values across all genes). We redirect the reader to Fig. 1A and Suppl. Info. A.1 for more details on the pipeline.

**Figure 1.**
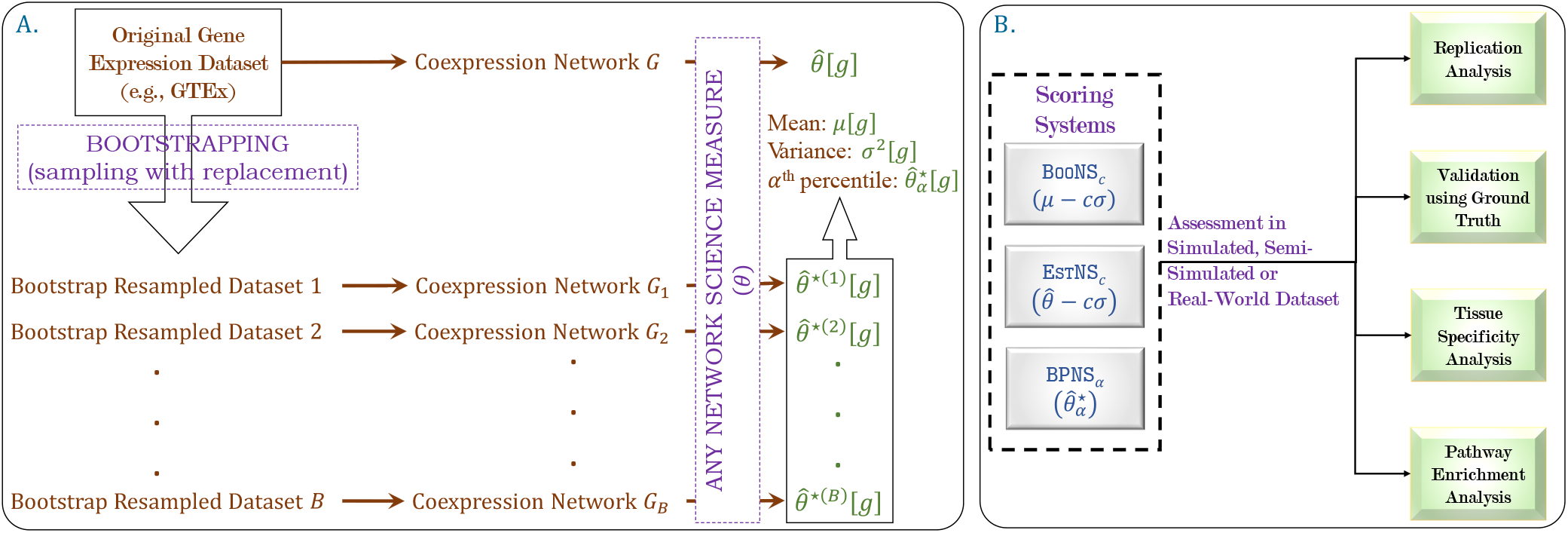
Our Study Overview. A. Our proposed framework to generate distribution of network science measurements with a focus on degree and PageRank centrality (although other network science measures can also be analyzed). B. Our proposed uncertainty-aware gene scoring systems along with different comparative assessments of gene rankings based on these scores across simulated, semi-simulated and real-world datasets, using different validation and biological interpretation approaches.

To highlight a stable central gene *g* with high *µ*[*g*] and low *σ*[*g*], we propose a simple uncertainty-aware gene score of the form *µ*[*g*] *−cσ*[*g*]. The vector, *µ −cσ*, comprising these scores of all genes, referred to as the `BooNS` scoring (for different values of *c*), induces a ranking of all genes.

The centrality score of the gene *g* in the coexpression network constructed on the original gene expression dataset with the actual number of samples, denoted as 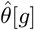 (Fig. 1A), is uncertainty-blind, and thus, we propose an uncertainty-aware gene score of the form 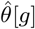 *− cσ*[*g*], calculated using the lower bound of the normal-approximation bootstrap confidence interval (10). Similar to that in `BooNS`, the corresponding vector, 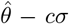, comprised of these scores across all genes, leads to another ranking, hereafter referred to as EstNS. It may be noted here that in both of these scorings *cσ* quantifies the gene-specific uncertainty in the measure; accordingly *c* is termed as the “*sigma-limit*” of the score. We use the notation `BooNS`_*c*_ and EstNS_*c*_ to denote the `BooNS` and EstNS scoring of the genes with sigma-limit *c* respectively.

Finally, we propose an additional uncertainty-aware score using the lower bound of the (1 *−* 2*α*) *×* 100% confidence interval of the bootstrap distribution for the gene *g*, i.e., the *α*^th^ percentile of the distribution, represented as 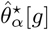. Ranking of the genes induced by the corresponding vector, 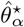, comprising these scores across all genes, is referred to as `BPNS`. Note that, unlike the previous two rankings wherein the centrality measurements are penalized based on summary statistics of the bootstrap distribution, this score penalizes uncertainty based on the shape of the distribution curve.

Hyperparameter tuning of the parameters *c* and *α* across all genes show minimal effect of their values on the gene rankings (Suppl. Info. A.2 and Suppl. Data S3). However, since larger sigma-limits and smaller percentiles are highly sensitive to outliers, we limit the values of *c* and *α* to *{*0, 1, 2*}* and *{*25, 49*}* respectively. It may be noted here that the scorings are uncertainty-blind when *c* = 0. We could’ve used other scorings based for instance on mean-variance analysis or risk-aware measures from modern portfolio management, but economists typically require a user-defined threshold to define these scorings (e.g., Sharpe and Sortino ratios (1), and exceedance probability (7)), which was against our preference for a simpler threshold-free score.

### 2.2. Analyses of Proposed Scorings

Considering the extensive use of centrality in downstream biological analysis (see Suppl. Info. A.3 and references cited therein), we provide a systematic assessment of the efficacy of our proposed scorings from the perspective of their validation rate, replicability, and tissue specificity analysis (Fig. 1B), relative to the baseline for degree and PageRank centralities. We use the uncertainty-blind scorings, i.e., EstNS_0_ and `BooNS`_0_ as baseline scores and compare performance of all uncertainty-aware scorings against them.

For validation and replication analyses (see Suppl. Info. A.4.1 and Suppl. Info. A.4.2 for detailed process of these analyses), we primarily analyze the percentage of overlapping genes (POG) (4) using correspondence at the top (CAT) (21) plots. A set of *k* genes is used as the ground truth or reference geneset, and POG @ *k* for any scoring *ψ* is computed as the size of overlap (intersection) between the top-*k* ranking genes by *ψ* and this reference geneset. We plot the POG @ *k* values (given in Suppl. Data S4) for all scorings across different values of *k* in these CAT plots. Naturally, rankings/scores that show high values of POG for different values of *k* (including for lower values of *k* corresponding to the top of the ranking) clearly recover the reference geneset better and should be preferred. To complement our CAT plots, we also provide the corresponding recall @ *k* plot (see Suppl. Info. A.5.1 for more details) for each CAT plot in Suppl. Data S5, and found that the performance trends observed in CAT vs. recall @ k plots are mostly similar. Each of these plots also show the average performance of a random ranking of the genes (denoted as the random scoring) to provide a baseline expectation (Suppl. Info. A.5.2).

In addition to conducting validation and replication analysis of our proposed scorings on the real-world (GTEx tissue) datasets, we also conduct tissue specificity analyses as detailed in Suppl. Info. A.4.3 and as summarized below. For each tissue of interest, we obtained genes that are highly expressed in the tissue based on the Human Protein Atlas resource (13; 35) (tissues absent in this resource are omitted from this analysis), and assembled these genes into a customized geneset for the tissue. These customized tissue-specific genesets are then used as the reference genesets in a Gene Set Enrichment Analysis (GSEA) (performed via WebGestalt (11)) to underscore scoring configurations that rank these tissue-specific genes higher than the other genes. For a given tissue and ranking of genes in the tissue by a scoring configuration, if the customized geneset for the same tissue is positively enriched in the ranking, then it is referred to as a “true positive”. All other positive enrichments are referred to as “false positive” enrichments. A proposed scoring configuration is of course preferable if the ranking it provides on different tissues results in more true positive and fewer false positive enrichments. Owing to their consistently better performance across validation and replication analyses, we chose to perform this analysis only for `BooNS`_1_, `BooNS`_2_, and `BPNS`_25_, and compared the results with respect to uncertainty-blind configurations, namely EstNS_0_ and `BooNS`_0_. Finally, we interpret the pathway enrichment results from `BooNS`_1_-based ranking (see Suppl. Info. A.4.4), owing to its consistently better performance as compared to the other scorings, as will be seen below. See Suppl. Info. A.5.3 for more detailed information on the implementation of the analyses described here.

### 2.3. Simulated/Semi-Simulated Datasets

We simulated a population dataset (sample size *s* = 500, 000) over two gene sets, *S*_1_ and *S*_2_, each containing 500 genes, under a bipartite network formulation. Briefly, expression values for genes in *S*_1_ were each generated independently from a standard normal distribution. Expression of each gene in *S*_2_ were then set to a linear combination of its neighboring genes in *S*_1_, where neighbors were chosen uniformly at random from the *S*_1_ genes (see Suppl. Info. A.6.1 and Suppl. Fig. S1A).

For each real-world (GTEx) tissue dataset of interest (see Section 2.4 and Table 1), we use it as a reference dataset to simulate a population dataset using the R package dependentsimr. This package uses a Gaussian copula approach to generate a simulated omics dataset that mimics the gene distributions and inter-dependencies in the reference dataset (38). See Suppl. Info. A.6.2 and Suppl. Fig. S2 for details.

**Table 1.** Tissues analyzed: *n* is number of protein-coding genes that are “selected” (as defined in Suppl. Info. A.6.3) for a tissue, and *s* is number of samples. Tissues chosen as representatives of a category and used majorly across analyses are shown in bold.

| (a) Category I |  |  | (b) Category II |  |  | (c) Category III |  |  |
| --- | --- | --- | --- | --- | --- | --- | --- | --- |
| Tissue | $n$ | $s$ | Tissue | $n$ | $s$ | Tissue | $n$ | $s$ |
| **Muscle Skeletal** | **15163** | **706** | Stomach | 16485 | 324 | Vagina | 16683 | 141 |
| **Whole Blood** | **14582** | **670** | Pancreas | 15848 | 305 | Brain Amygdala | 16414 | 129 |
| Skin Sun Exposed (Lower Leg) | 16646 | 605 | **Pituitary** | **17036** | **237** | Uterus | 16407 | 129 |
| Thyroid | 16607 | 574 | Brain Cerebellum | 16463 | 209 | Brain Substantia Nigra | 16413 | 114 |
| Lung | 16765 | 515 | Brain Cortex | 16536 | 205 | **Kidney Cortex** | **16450** | **73** |

The population degree/PageRank (*θ*) of the genes refer to their degree or PageRank centralities in the coexpression network inferred from a population dataset. We also subsampled the simulated and semi-simulated population datasets uniformly at random without replacement to obtain observed (discovery or replication) datasets (see Suppl. Fig. S1C and Suppl. Fig. S2); the entire bootstrapping process is then applied to each observed dataset to compute the corresponding coexpression networks and gene rankings. Generation of these discovery and replication datasets are detailed in Suppl. Info. A.4.1 and Suppl. Info. A.4.2.

### 2.4. Real-world (GTEx) Datasets

We computed the ranking of genes corresponding to each scoring (Suppl. Data S2) for 15 different tissues (Table 1) profiled across hundreds of individuals by the Genotype-Tissue Expression (GTEx) consortium (32). Specifically, we used the GTEx Release V8 dataset on all protein-coding genes, and particularly the single-tissue cis-expression Quantitative Trait Locus (eQTL) dataset downloaded on February 2025. These 15 tissues were chosen and categorized based on the number of samples denoted *s*: Category I with large *s*, Category II with moderate *s* and Category III with relatively fewer samples. Further, for tissue T from Category I, we subsampled the corresponding preprocessed dataset to generate two more datasets with sample sizes 237 and 73 (corresponding to sample sizes of “Pituitary”, a Category II tissue, and “Kidney Cortex”, a Category III tissue, respectively); T 237 and T 73 denotes these datasets. See Suppl. Info. A.6.3 and Suppl. Data S1 for details. Finally, we used the SRA dataset ‘SRP300916’ (34) (see Suppl. Info. A.4.2 and Suppl. Data S6) for Muscle Skeletal from the recount3 data repository (37) as the replication dataset exclusively for replication analysis.

## 3. Results

### 3.1. Results on Simulation Benchmarks

#### Setup and initial observations

We simulated a dataset with two set of genes (*S*_1_ and *S*_2_) under a bipartite formulation. As expected, we observe that increase in sample size leads to increased statistical power and hence more inferred edges (see Suppl. Fig. S1C). Additionally, we observe a non-monotonic relationship between *µ* and *σ*, with variability peaking at intermediate mean values. This suggests that uncertainty in degrees of the genes is highest in genes with intermediate degree centrality, rather than being uniformly distributed.

#### Validation with respect to population dataset

We first note that all the scorings show significantly high values of POG as compared to random for smaller values of *k* (see Section 2.2 for definition of POG calculated using the top-*k* ranking genes). POG values of all configurations across the different scoring systems follow a non-monotonic pattern – the values increases with increase in *k*, reaches close to 1 when *k* → 500 genes, following which a modest decrease and a subsequent increase in the values is observed. This observation is consistent with the nature of simulations as most of the genes in *S*_2_ would be ranked the highest.

For smaller values of *k*, i.e., when *k <* 500 genes, the eight configurations across all scoring systems that were analyzed segregate into two distinct groups – the group with the highest POG values comprising `BooNS` and `BPNS` scorings, and the group with lower POG values comprising all configurations of EstNS (Fig. 2A). While scorings within a group show very similar POG values, there is a noticeable gap in the performance between these groups, especially in observed datasets with larger sample sizes. Among scorings in the former group with the highest POG values, `BooNS`_2_ achieves the highest POG in several scenarios, and is followed closely by `BooNS`_1_ and `BPNS`_25_. For instance, in the observed dataset with 706 samples, `BooNS`_2_, `BooNS`_1_ and `BPNS`_25_ respectively show 0.78, 0.76, and 0.76 POG values when *k* = 100, and 0.96, 0.957, and 0.953 POG values when *k* = 300 (Suppl. Data S4).

**Figure 2.**
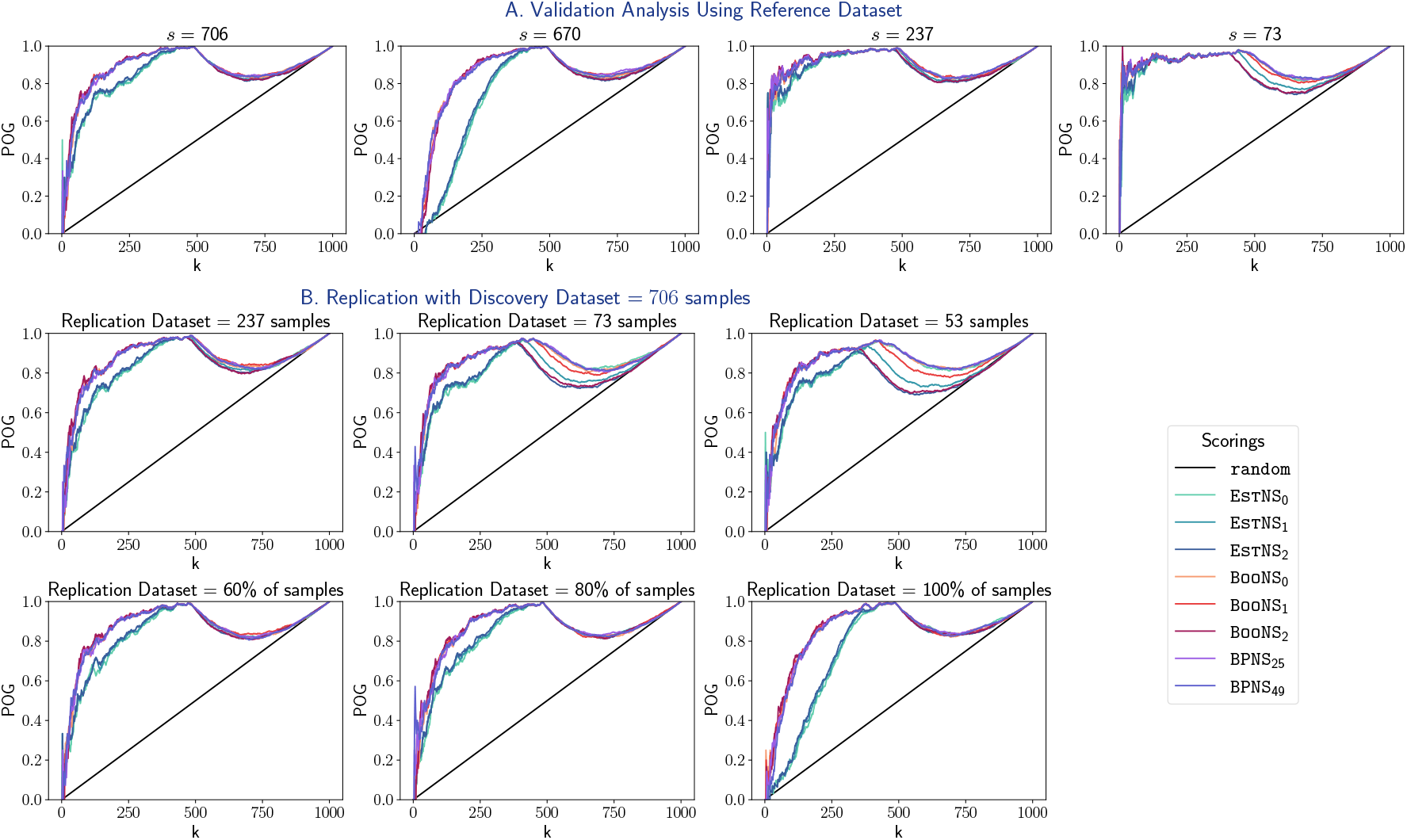
Analysis on Simulated Dataset for a Few Observed Datasets. A. CAT plots depicting the validation rate of all scorings relative to population for different observed datasets (with sample sizes matching that of certain real-world datasets). B. CAT plots depicting the replicability of all scorings in replication datasets with varied samples relative to a reference (discovery) dataset with 706 samples.

#### Replication between datasets

To assess replicability of the scorings across datasets, we used the observed datasets as discovery datasets, and corresponding to each discovery dataset, we sampled different replication datasets from the population with sample size at most the number of samples in the discovery dataset (see Suppl. Info. A.4.2). Finally, performance of all scores in the replication datasets were analyzed relative to their discovery dataset (Fig. 2B and Suppl. Fig. S3).

All the scorings show significantly high values of POG as compared to random for smaller values of *k* here as well. The non-monotonic pattern in the POG values followed by the scorings that is observed in the results for validation analysis is also evident here. Again, similar to that in validation analysis, for smaller values of *k*, i.e., when *k <* 500 genes, the scoring systems segregate into the same two distinct groups – `BooNS` and `BPNS` form the group with the highest POG values, and all configurations of EstNS form the group with lower POG values (Fig. 2B). Among `BooNS` and `BPNS`, `BooNS`_2_ achieves the highest POG in several scenarios, and is followed closely by `BooNS`_1_ and `BPNS`_25_. For instance, when the discovery and replication datasets have 706 and 53 samples respectively, `BooNS`_2_, `BPNS`_25_ and `BooNS`_1_ show the highest POG values of 0.84, 0.84, and 0.83 when *k* = 150 genes respectively (Suppl. Data S4).

#### Recommendations

Based on these results, we recommend the use of gene rankings based on the bootstrap distribution such as `BooNS`_2_, `BooNS`_1_ and `BPNS`_25_ for downstream analysis. However it remains to be seen (in the next sections) on whether real-world datasets characterized by heterogeneity in population, varied correlation structure among the genes, noise, etc., could benefit from such scorings.

### 3.2. Results on Semi-Simulated Dataset

#### Setup and initial observations

In accordance with the results on simulated datasets, we observe an increase in statistical power, and hence more inferred edges in the coexpression networks, with increase in sample size of the observed datasets (Suppl. Fig. S2). Additionally, there is negative correlation between *µ* and *σ* in observed datasets with larger sample sizes that slowly overturns into a positive correlation with decrease in sample size (Suppl. Fig. S2).

#### Validation with respect to population dataset

All the scorings show significantly high values of POG as compared to random, but, unlike in the simulated datasets, their POG values are highly similar to each other (Fig. 3A and Suppl. Fig. S4).

**Figure 3.**
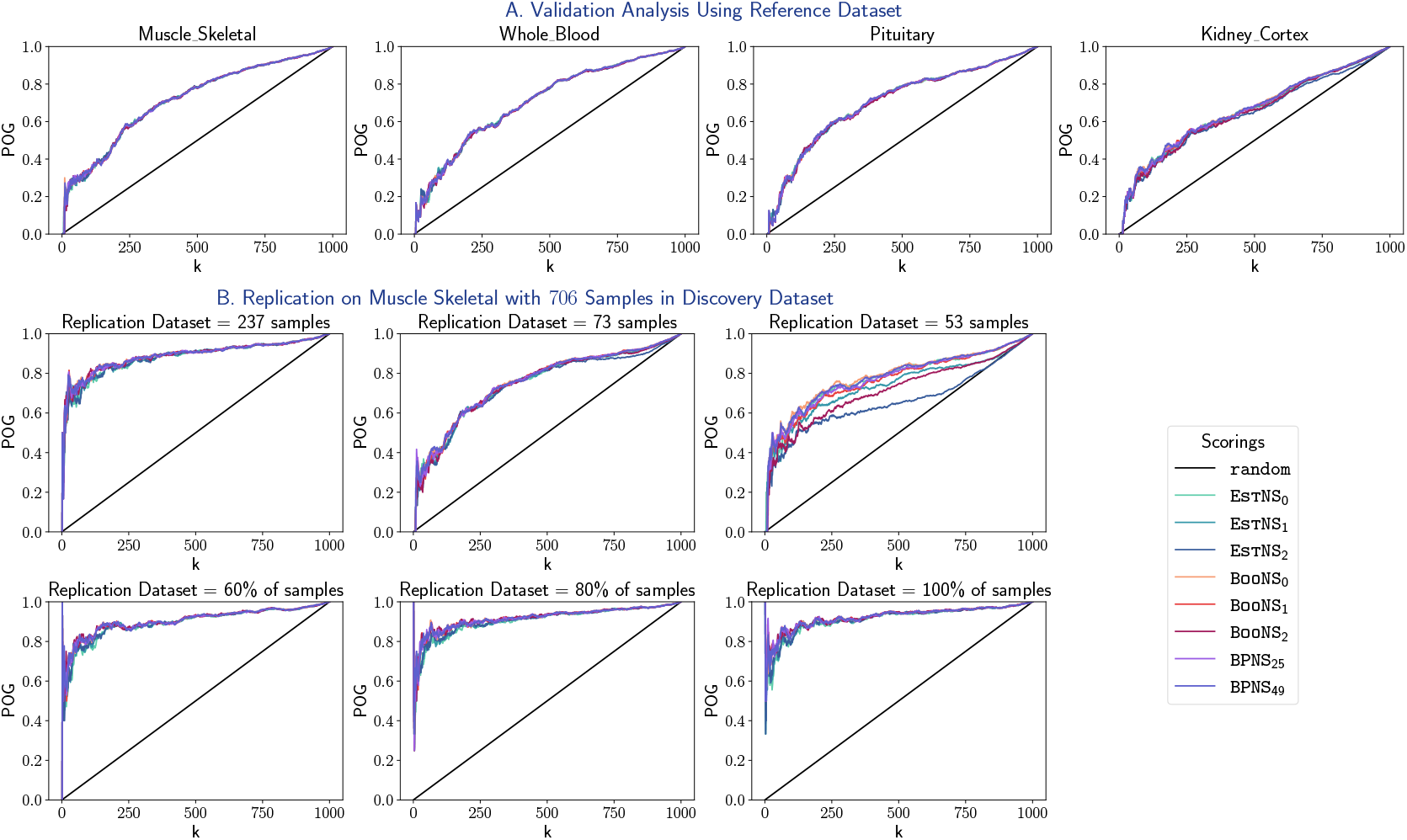
Analysis on Semi Simulated Dataset for a Few Tissues. A. CAT plots depicting the validation rate of all scorings relative to random scoring in observed datasets for a few tissues. B. CAT plots depicting the replicability of all scorings in replication datasets with varied samples relative to the observed dataset in Muscle Skeletal as reference (discovery) dataset.

#### Replication between datasets

Similar to the results previously observed in simulated datasets, all scorings show significantly high values of POG as compared to random. The scoring systems segregate themselves into the same two groups with similar POG values here as well; however, the average difference in POG between the two groups in these datasets is less as compared to that in simulated datasets. For instance, at *k* = 200, when both discovery and replication datasets of the Whole Blood tissue have the same sample size, i.e., *s* = 670, difference in POG values between `BooNS`_2_ and EstNS_1_ is close to 0.02 as compared to that of 0.265 in the simulated datasets (Suppl. Data S4). Additionally, we observe that `BooNS`_2_ and EstNS_2_ perform poorly under low sample size settings, such as when the replication dataset has at most 59 samples or the discovery dataset has 73 samples (pertaining to the Kidney Cortex tissue) (see Suppl. Fig. S5, Suppl. Fig. S6, and Suppl. Fig. S7).

#### Recommendations

The different rankings show similar performance in the validation analysis of semi-simulated data; so, the configurations recommended based on results in simulated data can also be followed here. Based on the replication analysis, we do not recommend `BooNS`_2_ and EstNS_2_ for low sample size settings.

### 3.3. Validation Analysis Relative to a Real-world Reference Dataset

To validate the stability of our scorings, we treat the preprocessed GTEx datasets comprising the original samples for Category I tissues as reference and their corresponding subsampled datasets with 237 and 73 samples as observed datasets. A directly proportional relationship between *µ* and *σ* is evident here (Fig. 4A and Suppl. Fig. S8).

**Figure 4.**
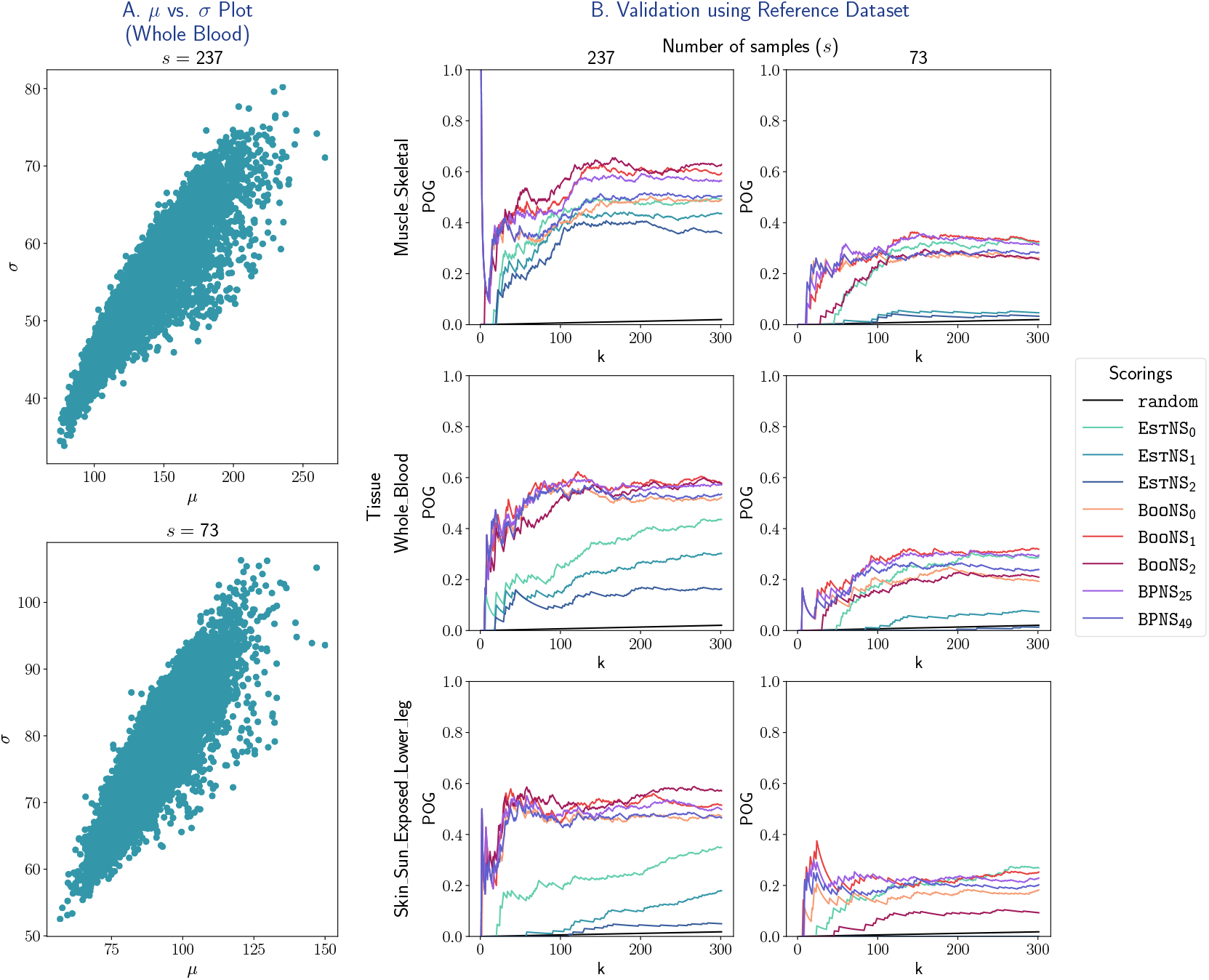
Validation of Degree Centrality Scorings using a Real-world Reference Dataset. A. Plots depicting the change in *σ* with respect to *µ*, and the effect of sample size *s* on the same for the Whole Blood tissue. B. CAT plots depicting the validation rate of all scorings in the subsampled datasets relative to the corresponding actual (preprocessed) dataset for Muscle-Skeletal, Whole Blood and Skin tissues.

#### Analysis of the CAT plots

We first note that several scoring systems show high POG values relative to that by random, whereas EstNS_1_ and EstNS_2_ exhibit poor performance consistently, even worse than random in several cases. Across all scenarios, `BooNS`_1_ and `BPNS`_25_ consistently rank among the top three in terms of their performance; `BooNS`_2_ has the highest validation rate on observed datasets with 237 sample size, but the performance of this scoring is subpar for datasets with 73 samples, possibly because it removes an excessive amount of variability (Fig. 4B and Suppl. Fig. S9). For instance, in analyses using the Muscle Skeletal tissue, `BooNS`_1_, `BooNS`_2_ and `BPNS`_25_ exhibit POG values at *k* = 100 genes of 0.48, 0.55, and 0.45 respectively when the observed dataset has 237 samples, but 0.28, 0.23, and 0.3 respectively when the observed dataset has 73 samples (Suppl. Data S4). Similarly, in the Lung tissue, when *k* = 150, these scorings have 0.44, 0.51, and 0.43 POG values respectively in datasets with 237 samples, but 0.25, 0.21, and 0.27 respectively in datasets with 73 samples (Suppl. Data S4). Also, as expected, the scorings exhibit higher POG @ *k* values when sample size is 237 as compared to when sample size is 73.

#### Recommendations

Clearly, validation rate of all the scorings increases with increase in number of samples, and hence we recommend the use of datasets with higher number of samples for downstream analysis. Also, given the promising results of `BooNS`_1_ and `BPNS`_25_, we recommend their application for network node centrality and other downstream analyses; however in large datasets with at least 237 samples, `BooNS`_2_ can also be preferred.

### 3.4. Testing Replication of GTEx Discovery Dataset Findings

In this section, we analyze the performance of all scorings in the replication dataset for Muscle Skeletal obtained from SRA (cf. Section 2.4) relative to that in the original GTEx dataset for the same tissue. We first note that the POG @ *k* values (Fig. 5) for all the scorings are very low as compared to that in the validation analysis, and can be attributed to dataset-specific variations like biological variability, noise, or differences in measurement conditions; however, all configurations of `BooNS` and `BPNS` exhibit higher POG than `random` for *k ≥* 818 which comprises only 5.60% of the genes (Suppl. Data S4). Conversely, all configuration of EstNS exhibit lower levels of POG than random for all *k <* 1000.

**Figure 5.**
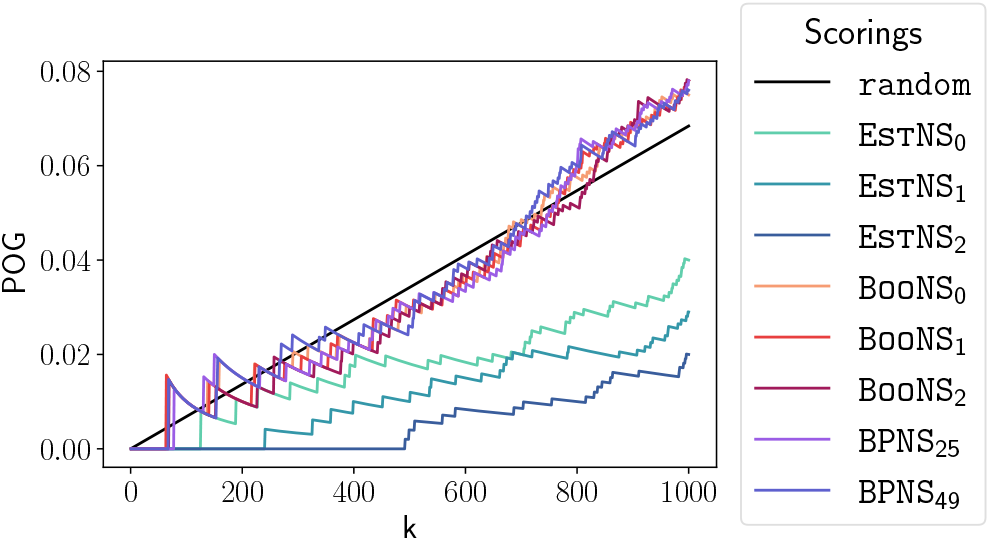
Replication of degree centrality scorings in real-world Muscle Skeletal datasets. CAT plot to assess replication of the scorings using discovery dataset as the reference and replication dataset as the query.

#### Recommendations

Although there is substantial effect of dataset-specific variations and the low sample size of the replication dataset on the results, all configurations of `BooNS` and `BPNS` show better replicability as compared to that by all configurations of EstNS, and hence should be preferred for further downstream analyses.

### 3.5. Results on Tissue Specificity Analysis

We used an external atlas dataset to assemble genes that are highly expressed in different tissues, and used these tissue-specific genesets to additionally validate our proposed scoring configurations. Specifically, we assessed if our scoring configurations applied on a tissue’s dataset resulted in significantly better ranks for the tissue-specific genes of that same tissue relative to other genes using GSEA enrichment analyses (reported at 5% and 10% FDR (False Discovery Rate) cutoffs; see Methods). Scoring configurations that exhibit this form of enrichment, also known as true positive enrichment, more often is preferred over configurations that lack any enrichment or exhibit false positive enrichments (i.e., enrichments for tissue-specific genes of a different tissue). See also Suppl. Fig. S10 for a visualization of these enrichments via boxplots of centrality scores of tissue-specific vs. other genes.

#### GSEA results

Heatmaps on the enrichment scores of the genesets (Fig. 6) clearly depict that EstNS_0_ performs poorly based on high false positive enrichments, whereas the other four rankings, namely `BPNS`_25_, `BooNS`_0_, `BooNS`_1_, and `BooNS`_2_, show very few false positives. We hence restrict further performance assessments to these four rankings only. For tissues in categories I and II, no significant differences were observed among the results by the configurations; notably, all of these configurations show true positive for every tissue in categories I and II except Whole Blood, and only 1 false positive is enriched in category I tissues. Additionally, for tissues in category III, in accordance with results from the previous analyses, `BooNS`_2_ shows comparatively poor performance (6 false positive enrichments) relative to the other configurations of `BooNS` (at most 3 false positives). The observations made here using an enrichment FDR cutoff of 5% shown in Fig. 6 largely hold when the FDR cutoff is relaxed to 10% (shown in Suppl. Fig. S11).

**Figure 6.**
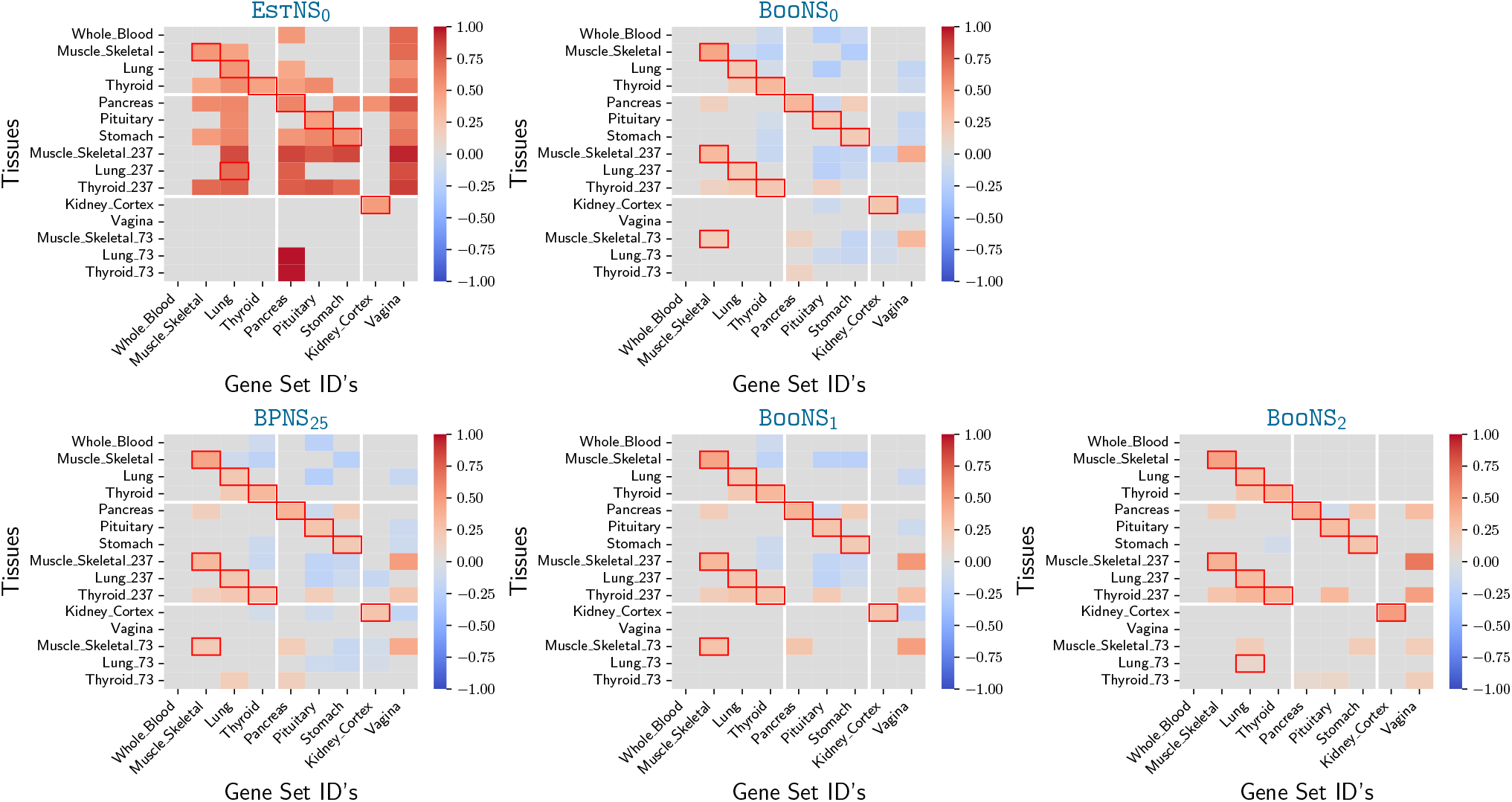
Tissue Specificity Analysis of Degree Centrality. Heatmaps showing the enrichment scores of the tissues, generated using WebGestalt. Only scorings that pass the 5% FDR threshold for enrichment have been highlighted; otherwise the enrichment score is manually set to zero. True positive enrichments has been marked using boxes.

#### Recommendations

For all tissues with at least 237 samples excepting Whole Blood, the uncertainty-aware rankings `BooNS`_1_, `BooNS`_2_, and `BPNS`_25_ perform significantly well. Based on this result, we recommend the use of datasets with at least 237 samples (as in category I and II tissues) when calculating centrality scores, and the use of uncertainty-aware rankings such as `BooNS`_1_ for tissue specificity analysis using GSEA. Similar to the analysis of semi-simulated datasets, this GSEA-based analysis also discourages the use of `BooNS`_2_ for low sample size settings.

### 3.6. Interpretation of Centrality Scores using Pathway Enrichment Analysis

We used the scoring by `BooNS`_1_ to analyze KEGG pathways that are significantly enriched across several or for a limited subset of tissues (Fig. 7). A total of about 262 pathways were significantly enriched across tissues (see “*Degree ES Table*.*xlsx*” in Suppl. Data S7), among which 49% (129) of the pathways were enriched in more than 7 tissues. Several of these pathways are known to be biologically significant, spanning across range of critical tissues; however, owing to this large number of enriched pathways, we only interpret a subset of them.

**Figure 7.**
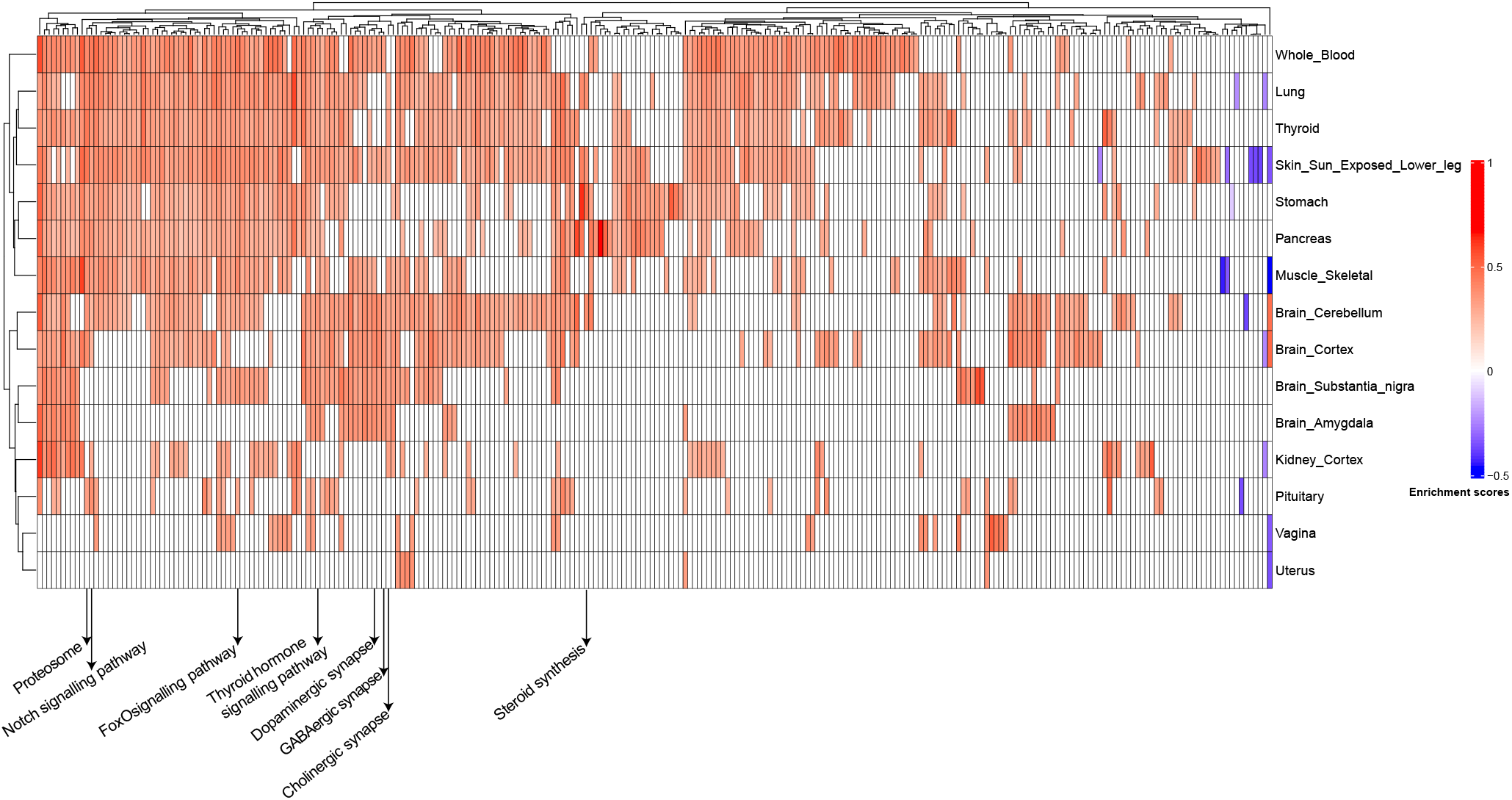
GSEA Analysis on KEGG Pathways. The heatmap shows the enrichment scores by `BooNS`_1_ scoring in different tissues, generated using WebGestalt. Only pathways that pass the 5% FDR cutoff on the adjusted p-values in a tissue are highlighted; otherwise the enrichment score is manually set to zero. To avoid visual cluttering, we annotate only the pathways that have been discussed in the manuscript.

#### Example pathways enriched across several tissues

The FoxO signaling pathway, which is involved in apoptosis, cell proliferation, and oxidative stress, shows strong associations with muscle skeletal via FoxO4, and with brain regions, kidneys, and lungs via FoxO3 (41). The Notch signaling pathway, a critical regulator of cell fate and tissue homeostasis, plays important roles in lungs, skeletal muscle, intestines, and brain regions (30). The thyroid hormone signaling pathway influences the development and metabolism of the brain, lungs, pituitary, thyroid and skeletal muscle (31).

#### Example pathways enriched for specific tissues

The GABAergic synapse pathway that plays a vital role in neurotransmission and overall function of the central nervous system (23), along with the Cholinergic and Dopaminergic synapse pathways, that contribute significantly to central nervous system regulation (27; 24), are predominantly enriched across several brain regions. Steroid biosynthesis, known to be involved in maintaining intestinal homeostasis (3), is strongly enriched in stomach. The proteasome pathway, connected to the ubiquitin–proteasome system (essential for muscle homeostasis) and whose dysregulation has been associated with muscle-related pathologies (22), is enriched in muscle skeletal.

#### Few examples of relevant top-5 genes in few pathways

(see “*Degree Main Table*.*csv*” in Suppl. Data S7) Consistent with prior studies showing that FoxO transcription factors are involved in regulating the *STK11* gene (25), this gene is among the top-5 genes in six out of the twelve tissues enriched for the FoxO signaling pathway. The gene product of the *APH1a* gene, which has a top ranking in six out of the nine tissues enriched for the Notch signalling pathway, is an important component of a protein complex in the same pathway (40). *SLC (Solute Carrier)* genes, which produce protein subunits that are involved in the synthesis of GABA (8), have top rankings in most of the brain regions. *SOAT1* gene, with a top ranking in most of the tissues enriched for the steroid biosynthesis pathway, is known to play a key role in cholesterol metabolism (17).

### 3.7. Performance of the Scorings for PageRank Centrality

Similar trends as observed in degree centrality also hold largely for PageRank centrality scores (see Suppl. Data S8 for similar plots on the gene rankings in PageRank centrality by all the scorings). But there are also minor deviations in trends as follows. First, genes with higher PageRank centrality measures are not seen with increase in sample size, unlike that in degree centrality (Fig. 4A), indicating that the newly detected edges (with increase in sample size) do not affect important paths and are possibly connected to already high degree influential nodes. Next, in validation analysis of GTEx category I tissues, there are certain scenarios such as low values of *k* when EstNS based PageRank centrality scorings perform better than other scorings (Suppl. Fig. S12), which is different from the trend observed for degree centrality (Fig. 4B). Finally, in tissue specificity analysis of category III tissues, the different `BooNS` configurations show similar number of false positive enrichments for PageRank centrality (3 to 4 false positives; Suppl. Fig. S13), unlike for degree centrality where `BooNS`_2_ exhibited a higher number of false positives (Fig. 6). Despite these few discrepancies across varied analyses, the overall performance trends remain the same, and the recommendations based on degree centrality are therefore also applicable for PageRank centrality.

## 4. Discussion

In this study, we were able to quantify the impact of uncertainty about the inferred coexpression network on downstream network analyses, in the context of degree and PageRank centrality of genes computed from several simulated, semi-simulated and real-world transcriptomic datasets. We achieve this primarily through our bootstrap-based uncertainty-aware `BooNS` and `BPNS` scorings, and their induced robust ranking of genes. We employed systematic validation and replication analyses using CAT plots, and tissue specificity analysis using GSEA to comparatively evaluate different rankings in their ability to recover reference or tissue-specific genes. A summary of key findings and network analysis recommendations from these analyses is distilled below.

- Prioritization of key genes is more robust using uncertainty-aware scorings from bootstrapped coexpression networks rather than centrality scores from a single network.
- Notably, our proposed scoring configurations such as `BooNS`_1_ and `BPNS`_25_ exhibit significantly stronger validation, replication and tissue specificity across the tested datasets, and hence recommended for network analyses over other configurations. Additionally, in large datasets with at least 237 samples, `BooNS`_2_ is also a viable approach.
- Based on our analyses of GTEx datasets, we recommend the use of transcriptomic datasets with roughly 250 or more samples for reliable network analyses.

Our comprehensive analyses between the datasets indicate a few inconsistencies in the results that can be attributed to dataset-specific variations. For instance, in replication analysis on real-world datasets, EstNS consistently shows poor performance than random, whereas in replication analysis on simulated and semi-simulated datasets on matched number of samples, such discrepancies are not observed. Similarly, the weak replication strength on real-world datasets can be attributed to biological variability and different measurement conditions. Another major challenge is the highly time-intensive nature to compute the coexpression networks and hence, the network science measures across a large number of bootstrap samples; consequently, this can be resolved with high-performance systems or time-efficient/parallel algorithms.

Despite the above caveats, our study demonstrates clearly that uncertainty about the coexpression networks indeed propagates to centrality measures, quantifies its negative impact on the same, and suggests a workaround in terms of uncertainty-aware gene rankings. Our systematic approach for quantifying and propagating network uncertainty using `BooNS` and `BPNS` can be broadly applied to study other network science measures as well in varied biological networks (such as those inferred from single-cell RNA-sequencing data that suffer from more technical noise than bulk transcriptomic data).

Bootstrap-based uncertainty estimation has been extensively used in phylogeny reconstruction (14), gene isoform expression quantitation (28), and gene regulatory network or Bayesian network reconstruction from genomic data (20; 16). However propagation of these uncertainty estimates to downstream analyses is not as widespread, and our work encourages active adoption of uncertainty-aware scorings and rankings to achieve the same.

## Supporting information

Supplementary Information (Supplementary Methods/Figures/DataFiles)

## Code and Data Availability

The code for bootstrap-based node scores’ generation and all other associated analyses is at https://github.com/BIRDSgroup/Bootstrap-based_Node_Scorings_BNS. The computed bootstrap-based node scores and all other associated results, for simulated, semi-simulated and real-world datasets, are publicly available at https://tinyurl.com/BNS-suppl-data (which aliases to https://drive.google.com/drive/folders/1Ru2zukLijqTYofn8lNLri2s7z9CTOS7i?usp=sharing).

## Acknowledgements

The authors thank Sanga Mitra for her help with the usage of WebGestalt. AI models such as ChatGPT and Gemini were utilized for copy-editing purposes (including polishing certain phrases or paragraphs) and minimal assistance with coding certain functions, and the Undermind AI platform was used to assist the literature review process. All AI-generated content were verified manually before inclusion in this work.

## Author Contributions

This work is performed as part of the doctoral thesis of SM with extensive inputs from MN. Implementation and analyses of the proposed metrics was performed by SM and interpretation of the pathway enrichment results was performed by NAS under the guidance of MN.

