## Supplementary Information (Supplementary Methods/Figures/DataFiles) for "Uncertainty-Aware Gene Rankings Reveal Key Players in Coexpression Networks"

, , , 1–21
Supplementary Information

### Uncertainty-Aware Gene Rankings Reveal Key Players in Coexpression Networks (Supplementary Information)

**Sugyani Mahapatra**[1,2,3] **Nilesh Anantha Subramanian**[1,2,3] **and Manikandan Narayanan**[1,2,3,4,*]

[1] Department of Computer Science & Engineering, Indian Institute of Technology (IIT) Madras, Chennai, India
[2] Centre for Integrative Biology and Systems medicinE, IIT Madras, Chennai, India
[3] Wadhwani School of Data Science & AI, IIT Madras, Chennai, India
[4] Sudha Gopalakrishnan Brain Centre, IIT Madras, Chennai, India

### A. Supplementary Methods

#### A.1. Description of our Workflow

We performed bootstrapping $1000$ times on the preprocessed (observed) gene expression dataset (the original GTEx datasets used across the analyses are provided in Suppl. Data S1) resulting in $1000$ bootstrap resampled datasets. $s$ samples were resampled uniformly at random with replacement from the original preprocessed dataset comprising of $s$ samples so as to generate a bootstrapped sample dataset with the same sample size. Each bootstrap resampled dataset resulted in a coexpression network $G_i$, and hence a measurement $\hat{\theta}^{\star(i)}$ for the genes; similarly, the original gene expression dataset also resulted in a coexpression network $G$, and hence a measurement $\hat{\theta}$ for the genes (see Suppl. Data S2 for these measurements on real-world datasets).

Traditional tools/packages like WGCNA (16), GWENA (17), CoGTEx (2) and GeneFriends (21) can be used for the construction of the coexpression network on the original preprocessed and on each of the bootstrap resampled datasets. However, in order to construct an undirected unweighted network without any assumptions on the degree of the genes, we preferred a simple network reconstruction using adjusted $p$-values. On a given gene expression dataset (original or bootstrap), the Spearman correlation coefficient and the related $p$-value between every pair of genes were calculated, following which the $p$-values were adjusted using Benjamini – Hochberg FDR correction. Pairs of genes which satisfied the 1% FDR cutoff were considered significantly correlated and edges were added between these genes.

The $1000$ coexpression networks on the bootstrap resampled datasets resulted in $1000$ measurements for each gene $g$; mean, standard deviation and $\alpha^{\text{th}}$ percentile of which are represented as $\mu[g]$, $\sigma[g]$ and $\hat{\theta}^{\star}_{\alpha}[g]$ respectively (with $\mu$, $\sigma$ and $\hat{\theta}^{\star}_{\alpha}$ referring to the vector of such values across all genes). Finally, scores/ranks of the genes (Suppl. Data S2) were computed using different scoring systems (Fig. 1B in main text).

#### A.2. Hyperparameter Tuning across Scoring Systems using Simulated and Semi-Simulated Datasets

We explored the effect of different hyperparameters across scoring systems on the gene ranks in both simulated (Suppl. Info. A.6.1) and semi-simulated datasets (Suppl. Info. A.6.2). For each of the scoring systems, we computed the rankings of all genes for several values of the corresponding hyperparameter ($c$ for EstNS and BooNS, and $\alpha$ for BPNS) and computed the Kendall's rank correlation coefficient of these ranks with the same in population (Suppl. Data S3); this was done for all simulated and semi-simulated datasets.

The maximum interquartile range (IQR) of the Kendall's $\tau$ values across most settings was found to be at most $0.09$, indicating low variability, thus, suggesting limited effect of these hyperparameters on the gene rankings when all genes are taken into consideration. However, a few exceptions were found wherein the IQR was as high as $0.2$, especially when the sample size of the dataset was $73$. Based on these observations, we set the values $\{0, 1, 2\}$ and $\{25, 49\}$ for $c$ and $\alpha$ respectively across all scoring systems and analyzed their performance in subsequent analysis.

#### A.3. Centrality in Identification of Hub Genes

Centrality based analysis have been extensively used in identification of essential components in a biological system (1; 14). While neighborhood-based or random-walk-based centrality measures like degree centrality, eigenvector centrality and PageRank centrality aid towards measuring the influence of a gene in the network, distance-based centrality measures like closeness centrality and betweenness centrality determine the communication and information transmission ability of the genes.

Degree centrality has been widely used in understanding the structural and functional characteristics of the system including but not limited to prediction of essential proteins and aging genes (30) and exploring the association between genes and complex diseases (31; 13). This centrality measure has also been extensively studied to validate (11; 20; 22; 8; 34; 23) and refute (37; 10; 24) the celebrated Centrality–Lethality hypothesis (12). On the other hand, PageRank centrality has played a pivotal role in capturing indirect connections and global network influence in biological systems (35; 15) and is highly instrumental in essentiality predictions (9; 36; 25) in multilayered (both weighted and unweighted) network systems.

Naturally, uncertainty in the prediction of centrality measures would lead to potential misinterpretations of the network systems (e.g., labeling a noisy unstable gene as highly central) affecting downstream analyses and decisions based on the same. Extensive use of centrality measures in different biological analyses, combined with the potentially detrimental effects of uncertainty in its prediction, makes it a good candidate for studying the propagation of uncertainty in biological networks.

#### A.4. Comparative Analyses of Proposed Scorings

##### A.4.1. Validation relative to a reference dataset

In validation analysis, we compare how correlated are the gene rankings by different scorings to the ground truth or the reference dataset. For simulated (Suppl. Info. A.6.1) and semi-simulated datasets (Suppl. Info. A.6.2), we used the top ranking genes, ranked by their population degree, as the reference set of genes.

However, with the real-world datasets, the underlying population is unavailable. Hence, we subsampled datasets with large number of samples (i.e., GTEx gene expression datasets of tissues belonging to Category I – see Suppl. Info. A.6.3, and Table 1 in main text) to generate observed datasets with smaller sample sizes, and analyzed the validation rate of gene rankings in these subsampled datasets relative to that by $\textsc{EstNS}_0$ i.e., $\hat{\theta}$ in the original preprocessed dataset with the actual number of samples. Thus, the reference geneset is computed as the top ranking genes by $\textsc{EstNS}_0$ in the coexpression network generated on the preprocessed gene expression dataset with the actual samples.

##### A.4.2. Replication across datasets

We also compared the stability in gene rankings by all scoring systems between two observed datasets. Alternatively stated, to analyze the replicability, we used two different observed sample datasets – the dataset with the larger number of samples acted as the discovery dataset and is used to compute the reference geneset, while the other with similar or fewer samples acted as the replication dataset. For every scoring $\psi$, top ranking genes with the highest centrality scores by $\psi$ in the discovery dataset was used as the reference geneset, and performance of $\psi$ in the replication dataset was analyzed relative to this geneset.

Observed datasets across varying sample sizes that were sampled from the simulated (Suppl. Info. A.6.1) and semi-simulated (Suppl. Info. A.6.2) population gene expression datasets acted as the discovery dataset. Corresponding to each such dataset, multiple replication datasets were separately sampled from the respective population with $60\%$, $80\%$ and $100\%$ sample size of the discovery dataset. Additionally, replication datasets with $237, 73$ and $53$ samples were also separately sampled from the respective population and were analyzed relative to the discovery datasets with $706$ sample size (i.e., the discovery dataset corresponding to the gene expression data for Muscle Skeletal tissue by GTEx (26)).

For replication analysis on real-world datasets, we used the preprocessed gene expression dataset for Muscle Skeletal tissue from the GTEx data repository (see Suppl. Info. A.6.3) comprising of the original $706$ samples as the discovery dataset and the gene expression dataset for the same tissue on $53$ healthy individuals corresponding to the SRA project id 'SRP300916' (GEO accession number 'GSE164471' and BioProject ID 'PRJNA690800') (27) from the recount3 data repository (32) as the replication dataset (see Suppl. Data S6). This replication dataset originates from the Sequence Read Archive (SRA) and was processed following the recount3 pipeline. However, since the replication dataset consists of read counts instead of the preprocessed eQTL data, unlike the discovery dataset, we performed Trimmed Mean of M-values (TMM) Normalization and Inverse Normal Transformation (in accordance with the gene expression normalization techniques used by the GTEx repository) for the preprocessing pipeline. Thereafter, the data was adjusted for age and gender covariates as provided with the project. Following this, gene rankings by the different scorings were computed only on the $14619$ protein-coding genes that are common to both the datasets.

##### A.4.3. Tissue specificity analysis

Genes with elevated expression levels in a tissue, hereafter, known as "tissue-specific genes", are highly involved in several functionalities of the tissue and should be highly central in the underlying coexpression network for the corresponding tissue. The Human Protein Atlas (5; 28) categorizes tissue-specific genes (also known as "tissue elevated genes") into "*Tissue Enriched*", "*Group Enriched*" and "*Tissue Enhanced*". In this analysis, we assess how tissue-specific are the genes with top ranks by the different scorings. Since the gene expression values in simulated and semi-simulated datasets are randomly generated, this analysis is performed only on GTEx datasets.

Tissue-elevated genes ('total elevated' category) for Kidney Cortex (7), Lung (18), Muscle Skeletal (19), Pancreas (3), Pituitary (3), Stomach (6), Thyroid and Vagina and Cell-type elevated genes for Blood and Immune Cells (henceforth termed as Whole Blood) (29) were downloaded from the Human Protein Atlas (5; 28) and used as the tissue-specific geneset for the respective tissues. All these genesets were assembled into a customized "ground truth" reference geneset database. Enrichment score of these genesets by the different scorings in each of these tissues were computed using WebGestalt (4) (the `WebGestaltR` package (version 0.4.6) for the R programming language version 4.5.2 was used in the implementation of the same). Only enrichment scores that pass the 5% FDR cutoff on the adjusted $p$-values are determined to be significant and hence, are analyzed.

Each of the scorings that were used to perform this analysis induce different rankings of the genes; clearly, scores that positively enrich the gene set corresponding to the specific genes for the same tissue are preferable over others. We say the positive enrichment of a geneset in any tissue is a "true positive" if it corresponds to the specific genes of the same tissue; otherwise we say the enrichment is a "false positive".

##### A.4.4. Interpretation using GSEA

We used the centrality measures by $\texttt{BooNS}_1$ to perform GSEA on the KEGG pathway database and analyzed significantly enriched pathways using WebGestalt (4). Similar to the tissue specificity analysis, only pathways with enrichment scores that pass the 5% FDR cutoff on the adjusted $p$-values were considered "significantly enriched" for the corresponding tissue and analyzed further; enrichment scores for all other pathways were manually set to zero. Union of enriched pathways across all tissues (generated from the GSEA) were assembled into an enrichment matrix (Suppl. Data S7) using their enrichment scores, and depicted as a clustered heatmap for visualization and interpretation. Owing to the promising results by $\texttt{BooNS}_1$, this analysis was performed on the gene rankings by this scoring only so as to assess their involvement and contribution in determining significant functional pathways and in elucidating associated biological processes.

#### A.5. Techniques used across Analyses

##### A.5.1. Computation of recall @ $k$ plots

Similar to the CAT plots, the recall @ $k$ plots were also generated by plotting the recall values of the scorings with respect to a reference geneset for different values of $k$. However, the reference geneset used here was fixed apriori and is independent of $k$ – we used a set of $1000$ genes as the reference (needs clarifications). The recall @ $k$ value for any scoring $\psi$ is computed as the size of overlap (intersection) between the top-$k$ ranking genes by $\psi$ and this reference set of genes.

##### A.5.2. The `random`-ranking

In both the CAT and recall @ $k$ plots, we also plotted the expected values of POG/recall @ $k$ in a random assignment of ranks to the genes as a separate score, termed as `random`. This scoring system assigns a rank to each of the $n$ genes in a gene expression dataset uniformly at random. Alternatively stated, out of all possible $n!$ rankings of the genes, one of them is picked uniformly at random.

The POG/recall @ $k$ of this score is also computed with respect to a reference gene set, denoted as $R$. Let the set of top-$k$ ranking genes from the chosen random assignment be represented as $A$. Stated differently, $A$ represents a set of genes constructed by picking $k$ genes from the input geneset uniformly at random without replacement; we say a "pick" to be a "success" if the chosen gene $g \in R$, and "failure" otherwise. Then the random variable $X = |A \cap R|$ follows a hypergeometric distribution with the expectation $\mathbb{E}[X] = \frac{k|R|}{n}$. Thus, the expected value for both POG @ $k$ and recall @ $k$ can be computed as $\frac{\mathbb{E}[X]}{|R|} = \frac{k}{n}$.

##### A.5.3. Implementation details

All analyses on simulated and semi-simulated datasets and preprocessing of the real-world GTEx datasets were performed on an Intel Core i7 processor running at 1.30 – 1.50 GHz using 8 GB memory with Windows 11 (Home) (64 bit operating system and x64-based processor). While all simulations and analyses of the synthetic datasets were performed using the Python version 3.12.5 programming language, preprocessing of the real-world GTEx datasets were performed using the R version 4.4.2 programming language with parallel processing on a cluster of $5$ cores to ensure faster covariate analysis.

The bootstrapping process, coexpression network construction and centrality computation for the real-world datasets were performed on a local server with Intel Xeon Platinum 8180 CPU running at 2.5 GHz frequency with 56 physical cores comprising of two threads per core and using 1 TB memory with CentOS 7 operating system. The programs were executed parallelly on $50$ cores using Python version 3.10 programming language. The internal threads `OMP_NUM_THREADS`, `MKL_NUM_THREADS`, `NUMEXPR_NUM_THREADS`, `OPENBLAS_NUM_THREADS` and `VECLIB_MAXIMUM_THREADS` were set to $1$ during the bootstrapping and coexpression network process and to $10$ for the centrality measures computation. Random sampling of the individuals for bootstrapping was performed using the random package in Python version 3.10 and the seeds used have been recorded. We used the `full_matmul_symmetrical` and `derive_pvalues` functions from the `corals.correlation` package for the correlation coefficient and p-value computation respectively and the `multipletests` function from the `statsmodels.stats.multitest` package for the FDR correction. All related programs were merged together using the Bourne Again SHell (BASH) scripting language.

Pathway enrichment analysis (GSEA) was performed using the WebGestaltR package (version 0.4.6) (4) in the R programming language version 4.2.1, which provides an interface to the WebGestalt web server; GSEA analysis was performed on the KEGG pathway database with 'gene symbols' as the type of gene input. The analysis was performed on an Intel Core i7 processor running at 3.20 GHz using 16 GB memory with Windows 11 Pro (64-bit operating system and x64-based processor).

#### A.6. Datasets used across Analyses

##### A.6.1. The simulated dataset

We simulated a gene expression dataset for a population with $1000$ genes and five hundred thousand samples. The gene expression values for the first set of $500$ genes $S_1$ (comprising of $g_1$ to $g_{500}$) were each independently generated by sampling from a standard normal distribution. For each gene $g$ in the remaining set of $500$ genes $S_2$ (comprising of $g_{501}$ to $g_{1000}$), its neighbors were chosen by first choosing its degree uniformly at random from $1$ to $500$ and then randomly choosing that many neighbors from $S_1$ (see Suppl. Fig. S1A).

Following this, the gene expression values for $g \in S_2$, represented as the column vector $E_g$, were sampled as a linear function of the gene expression values of its corresponding neighbors in $S_1$ as follows. Let the matrix $\mathbf{D} \in \mathbb{R}^{500000\times 500}$ be the gene expression dataset containing the expression values of the genes in $S_1$. Then, $E_g$ is calculated as:

$$E_g = \mathbf{D}\vec{\alpha} + 0.1\,\vec{\mathbf{1}} + 0.01\,\vec{\epsilon}$$

Here, the noise vector $\epsilon$ is independently sampled from a standard normal distribution and $\vec{\alpha}$ is initialized as:

$$\alpha_i = \begin{cases} 0.995, & \text{if } g_i \in S_1 \text{ is a neighbor of } g \\ 0, & otherwise \end{cases}$$

The population degree/PageRank ($\theta$) of the genes refer to their degree or PageRank centrality measures in the coexpression network constructed on this (population) dataset. Suppl. Fig. S1B shows the adjacency matrix of this coexpression network (represented as a heatmap) and the population degree distribution of the genes. From this population, four observed gene expression datasets with sample sizes corresponding to certain tissues in the real-world datasets (cf. bold-faced tissues in Table 1 in main text) were constructed by sampling uniformly at random without replacement, and the entire bootstrapping process was applied to compute the corresponding coexpression networks and gene rankings.

##### A.6.2. The semi-simulated dataset

For each real-world (GTEx) tissue dataset of interest (see Suppl. Info. A.6.3, and Table 1 in main text), we use it as a reference dataset to simulate a population dataset using the R package `dependentsimr`. A Gaussian copula approach is used by `dependentsimr` to generate a simulated omics dataset that mimics the gene distributions and inter-dependencies in the reference dataset (33).

In detail, we first extracted gene expression values of the top-$1000$ protein-coding genes with the highest variance across individuals in the original gene expression dataset of the tissue. Dependency structure among genes in this data was then inferred using `dependentsimr`'s `get_random_structure` function. We call this function with appropriate parameters: "`normal`" to simulate approximately normally distributed expression value; and "`corp.cor`" to preserve inter-gene correlation structure, thereby preserving biologically relevant co-expression patterns seen in the reference dependency structure. Subsequently, $500,000$ random samples were drawn based on this structure to constitute a population gene expression dataset over the chosen $1000$ genes. Using this semi-simulated population dataset, we construct a coexpression network and compute gene centralities to obtain the population degree/PageRank ($\theta$) of genes in the tissue.

We subsequently sample the semi-simulated population dataset uniformly at random without replacement to obtain an observed (discovery or replication) dataset for the tissue; the entire bootstrapping process is then applied to the observed dataset to compute the corresponding coexpression networks and gene rankings. More details on the discovery and replication datasets are in Suppl. Info. A.4, and Suppl. Fig. S2.

##### A.6.3. Real-world (GTEx) datasets

We computed the ranking of genes corresponding to each scoring (see Suppl. Data S2) for $15$ different tissues profiled across hundreds of individuals by the Genotype-Tissue Expression (GTEx) consortium (26). Specifically, GTEx Release V8 dataset on all protein-coding genes, and particularly the single-tissue cis-expression Quantitative Trait Locus (eQTL) dataset downloaded on February 2025, was used. These $15$ tissues were chosen and categorized based on the number of samples denoted $s$: Category I with large $s$, Category II with moderate $s$ and Category III with relatively fewer samples. See Table 1 in main text for a list of all considered tissues and the number $n$ of "selected" genes in each tissue. A gene is selected for inclusion in the coexpression network if its expression is at least $0.1$ Transcripts Per Million (TPM) and it has $6$ mapped reads in at least 20% of the samples as analyzed by GTEx (these criteria are similar to that followed in GTEx's cis-eQTL dataset used here (26)).

The datasets were preprocessed to retain only protein-coding genes and adjusted for potential confounding covariates (Suppl. Data S1; computed centrality values are in Suppl. Data S2). For the covariate adjustment, we used all available covariates for the respective tissue, which includes top $5$ genotyping principal components, several inferred covariates (identified using the Probabilistic Estimation of Expression Residuals (PEER) method), sequencing platform, sequencing protocol and gender (again these preprocessing choices follow the analyses employed in GTEx cis-eQTL data (26)).

Apart from these, we subsampled the preprocessed datasets of Category I tissues to generate two more datasets with sample sizes $237$ and $73$ (corresponding to the number of samples in "Pituitary", a Category II tissue, and "Kidney Cortex", a Category III tissue). All of these subsampled datasets were generated by picking the samples in the actual preprocessed datasets uniformly at random without replacement.

Finally, we also used the SRA dataset 'SRP300916' (27) (see Suppl. Info. A.4.2 and Suppl. Data S6) for Muscle Skeletal from the recount3 data repository (32) as the replication dataset exclusively for replication analysis.

### B. Supplementary Figures

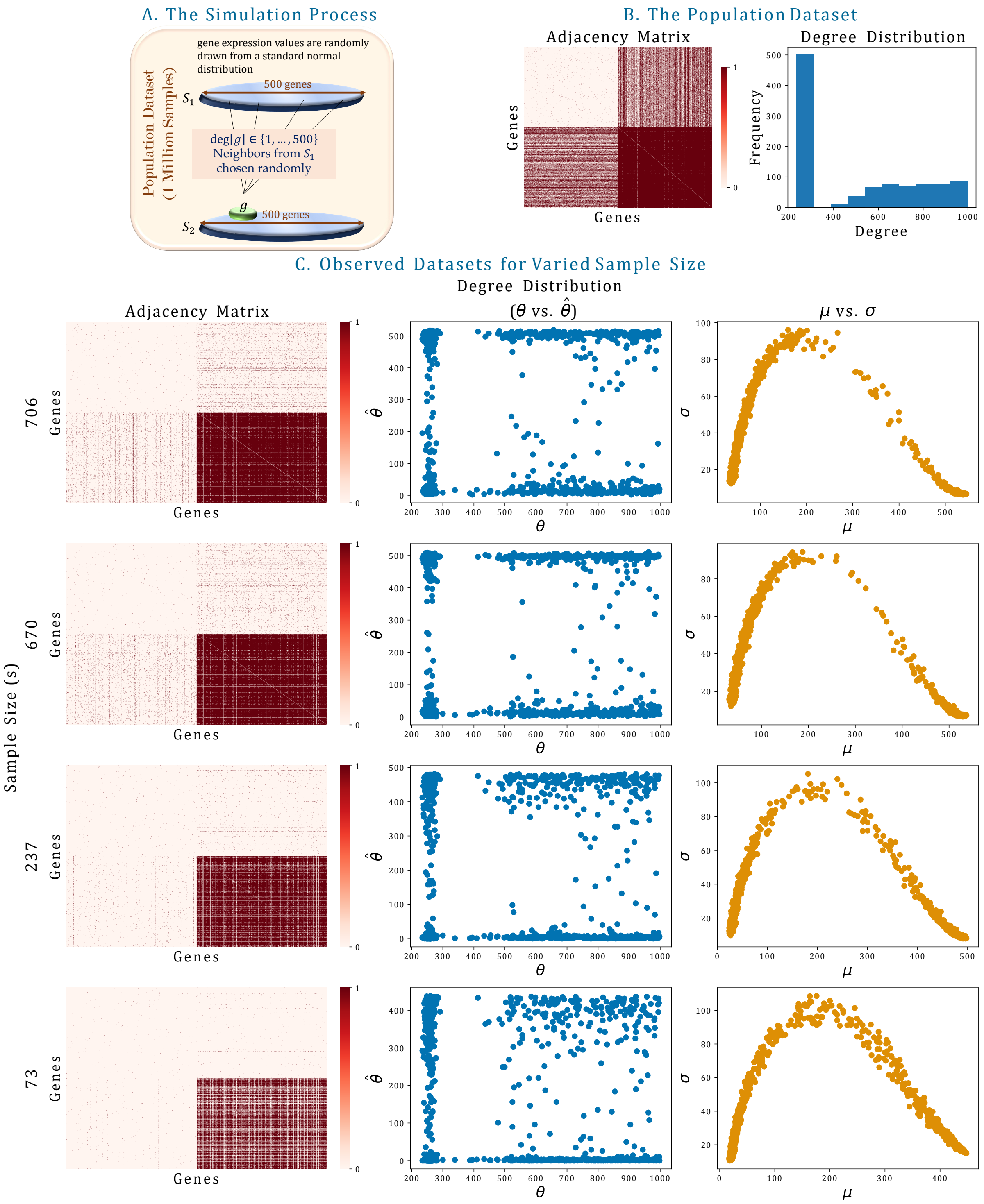


**Suppl. Fig. S1 Simulations.** A. The process of simulating the population dataset is briefly illustrated. B. Adjacency matrix of the underlying coexpression network represented as a heatmap and degree distribution of all genes in the simulated population. C. Similar adjacency matrices as in B., along with scatter plots depicting degree distribution of the genes with respect to their population degrees, and scatter plots depicting the change in $\sigma$ with respect to $\mu$ in a few observed datasets. Note that while genes in $S_1$ are mostly connected to genes in $S_2$ only, there are a lot of intra-set edges among the genes in $S_2$; this is possibly due to the linear dependencies among these genes introduced by the simulation design.

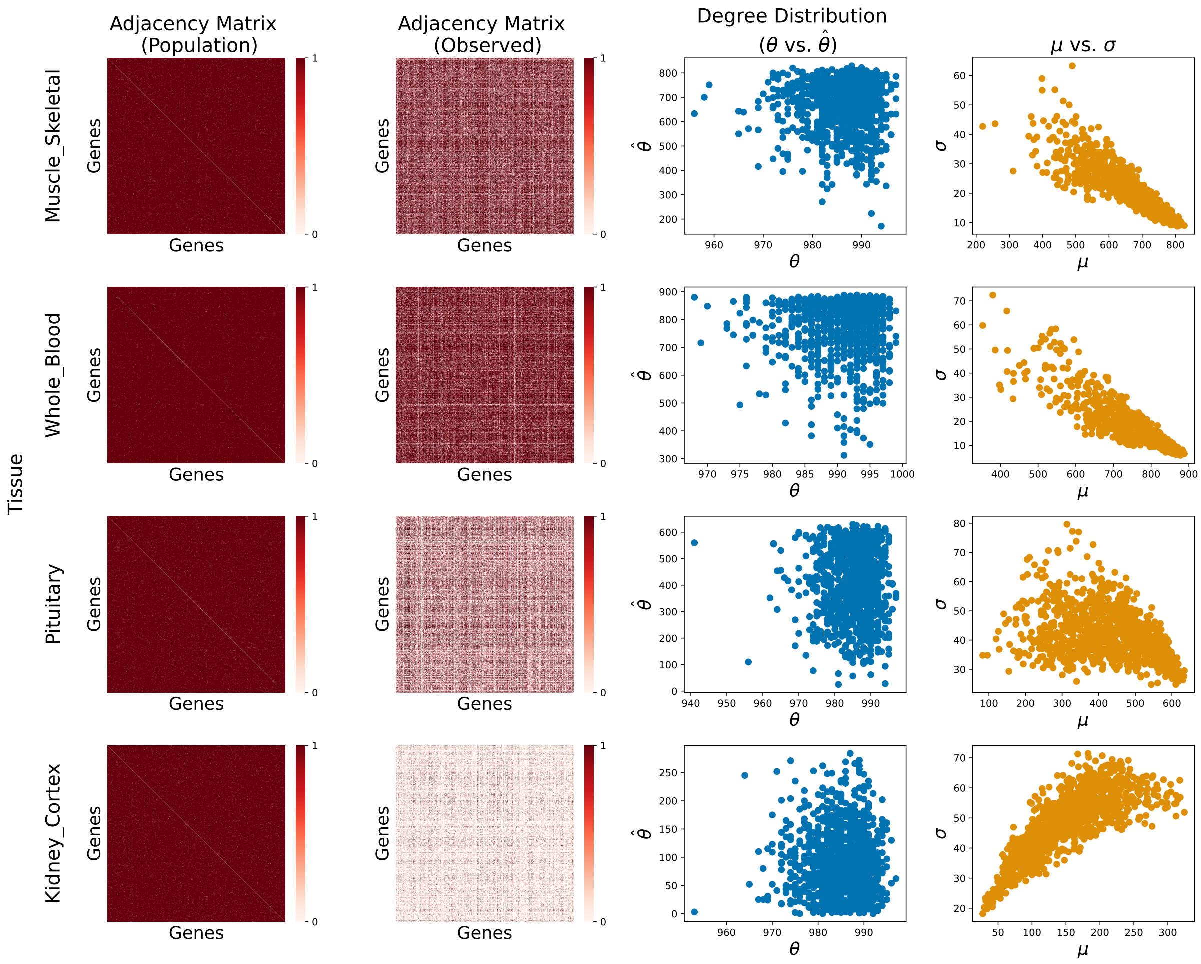


**Suppl. Fig. S2 Semi-Simulated Datasets.** Adjacency matrices of the coexpression networks (represented as heatmaps) of the simulated population and sampled observed datasets, along with scatter plots depicting estimated degree distribution of the genes in observed dataset with respect to their population degrees, and scatter plots depicting the change in $\sigma$ with respect to $\mu$ for Muscle Skeletal ($s = 706$), Whole Blood ($s = 670$), Pituitary ($s = 237$) and Kidney Cortex ($s = 73$) tissues.

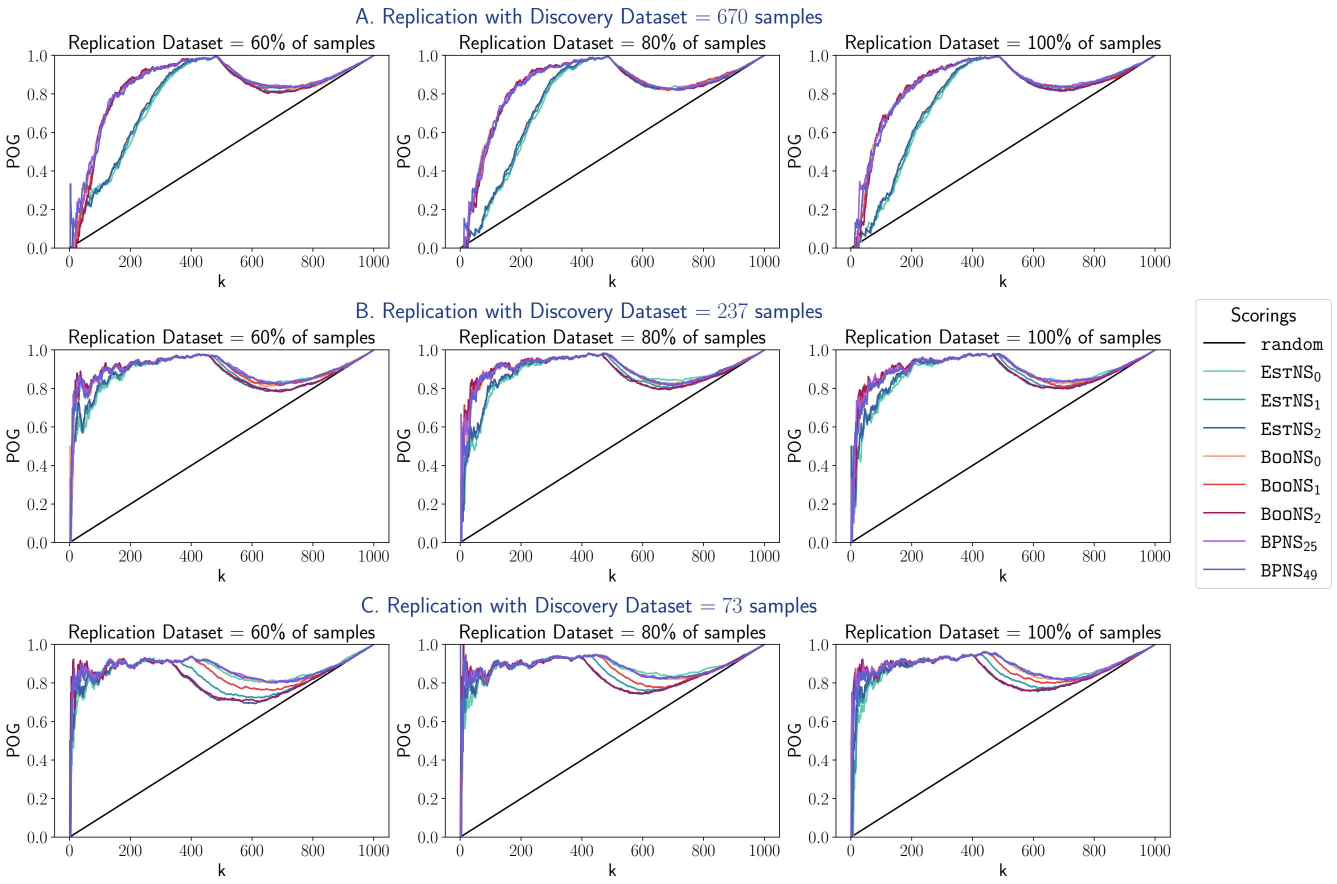


**Suppl. Fig. S3 Replication Analysis on Simulated Dataset.** CAT plots depicting the replicability of all scorings relative to different reference observed (discovery) datasets with varied samples. The plots are shown for replication datasets with different sample sizes relative to that of the discovery dataset, with the POG values for random assignment of ranks as a dashed line.

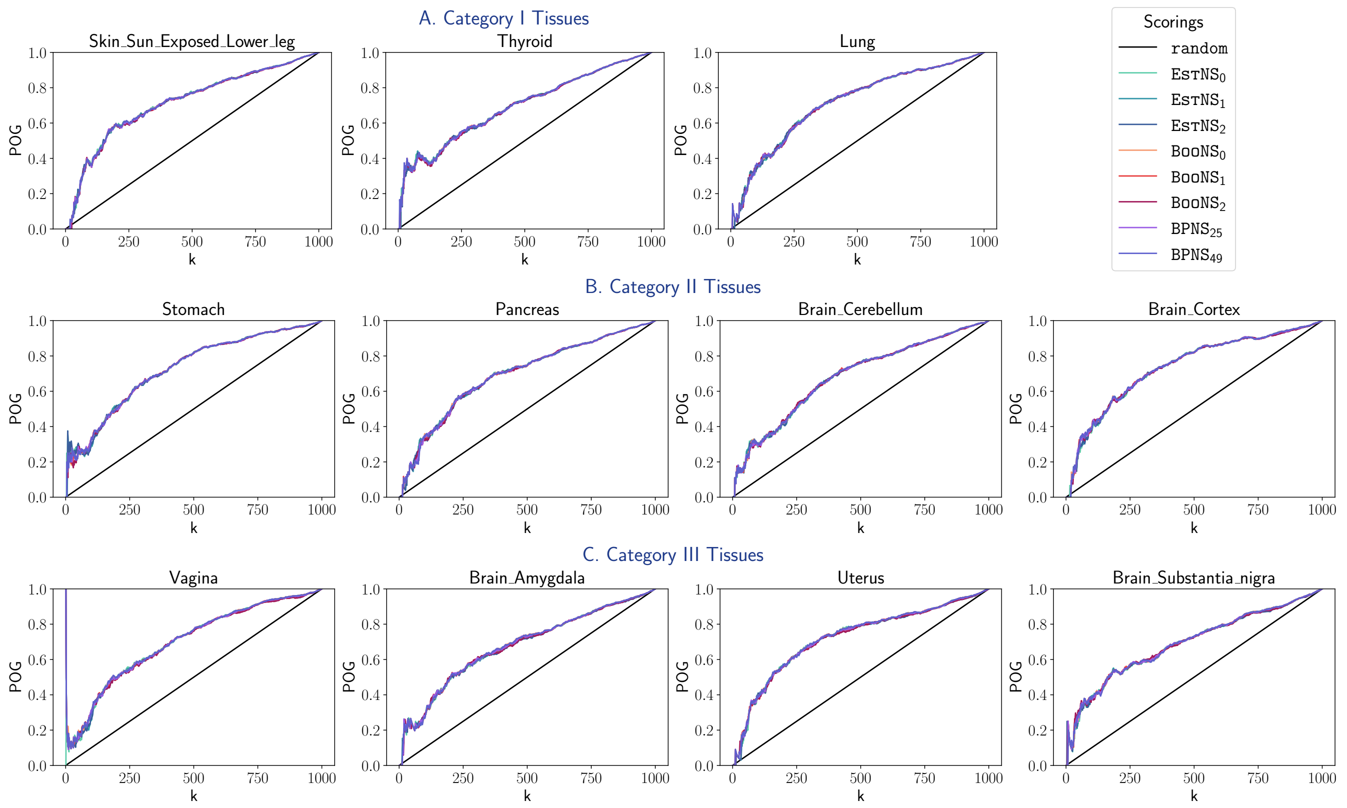


**Suppl. Fig. S4 Validation in Semi Simulated Dataset of Other Tissues.** Similar CAT plots as in Fig. 3A depicting the validation rate of all scorings.

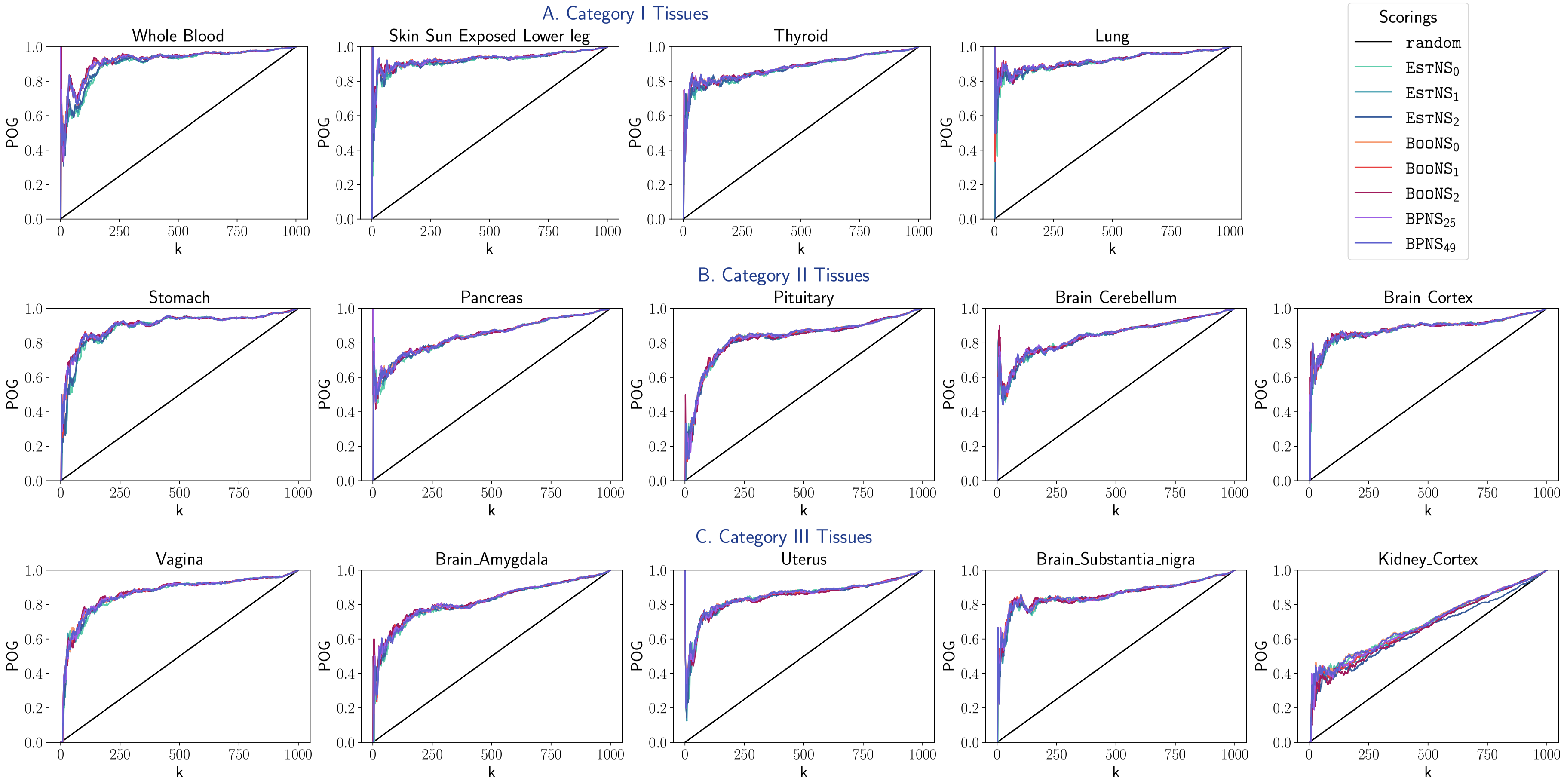


**Suppl. Fig. S5 Replication Analysis in Semi Simulated Dataset of Other Tissues** with replication dataset size $= 100\%$ sample size of discovery dataset. Similar CAT plots as in Fig. 3B depicting replicability of all scorings.

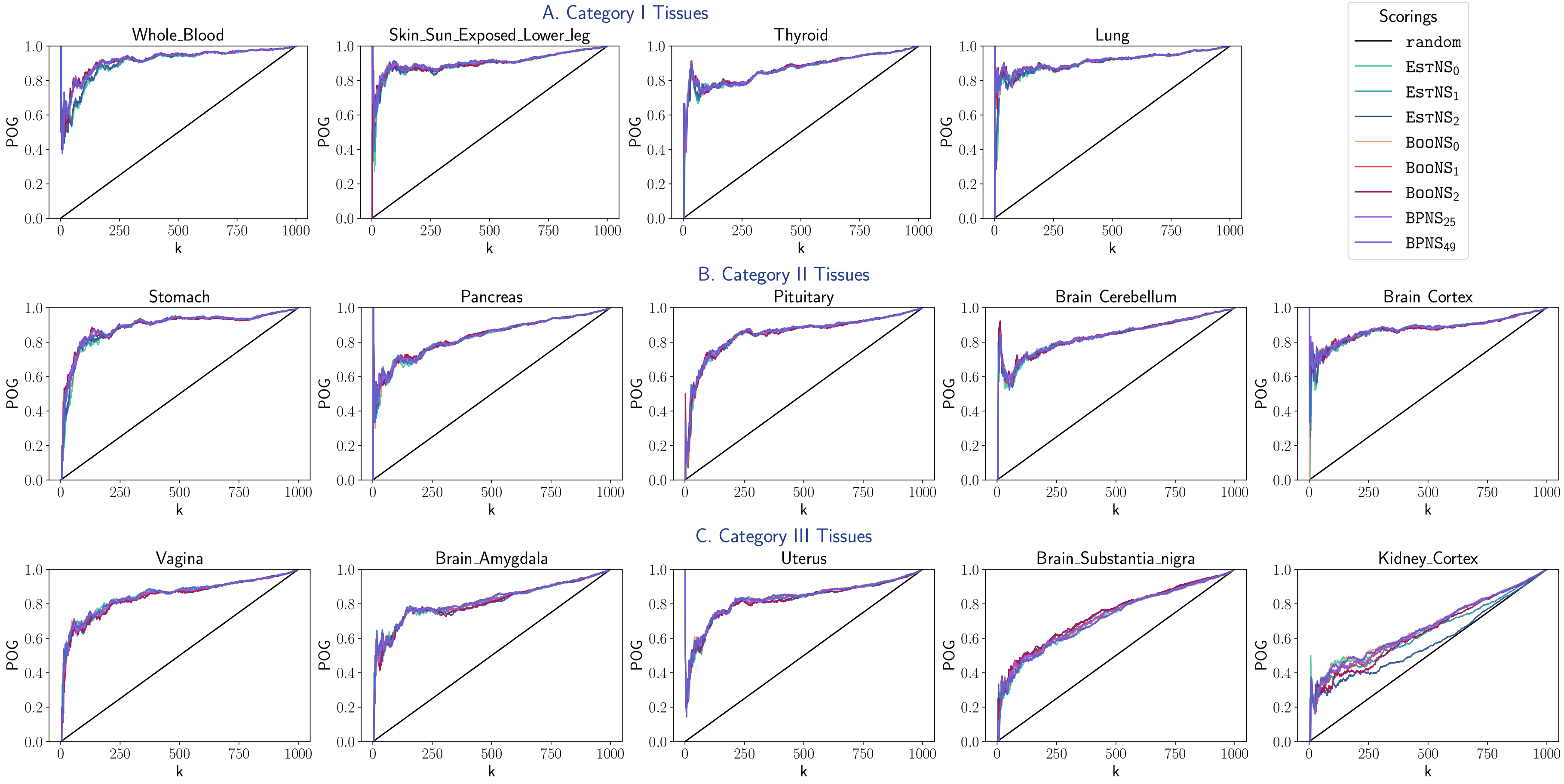


**Suppl. Fig. S6 Replication Analysis in Semi Simulated Dataset of Other Tissues** with replication dataset size = 80% sample size of discovery dataset. Similar CAT plots as in Suppl. Fig. S5 depicting replicability of all scorings.

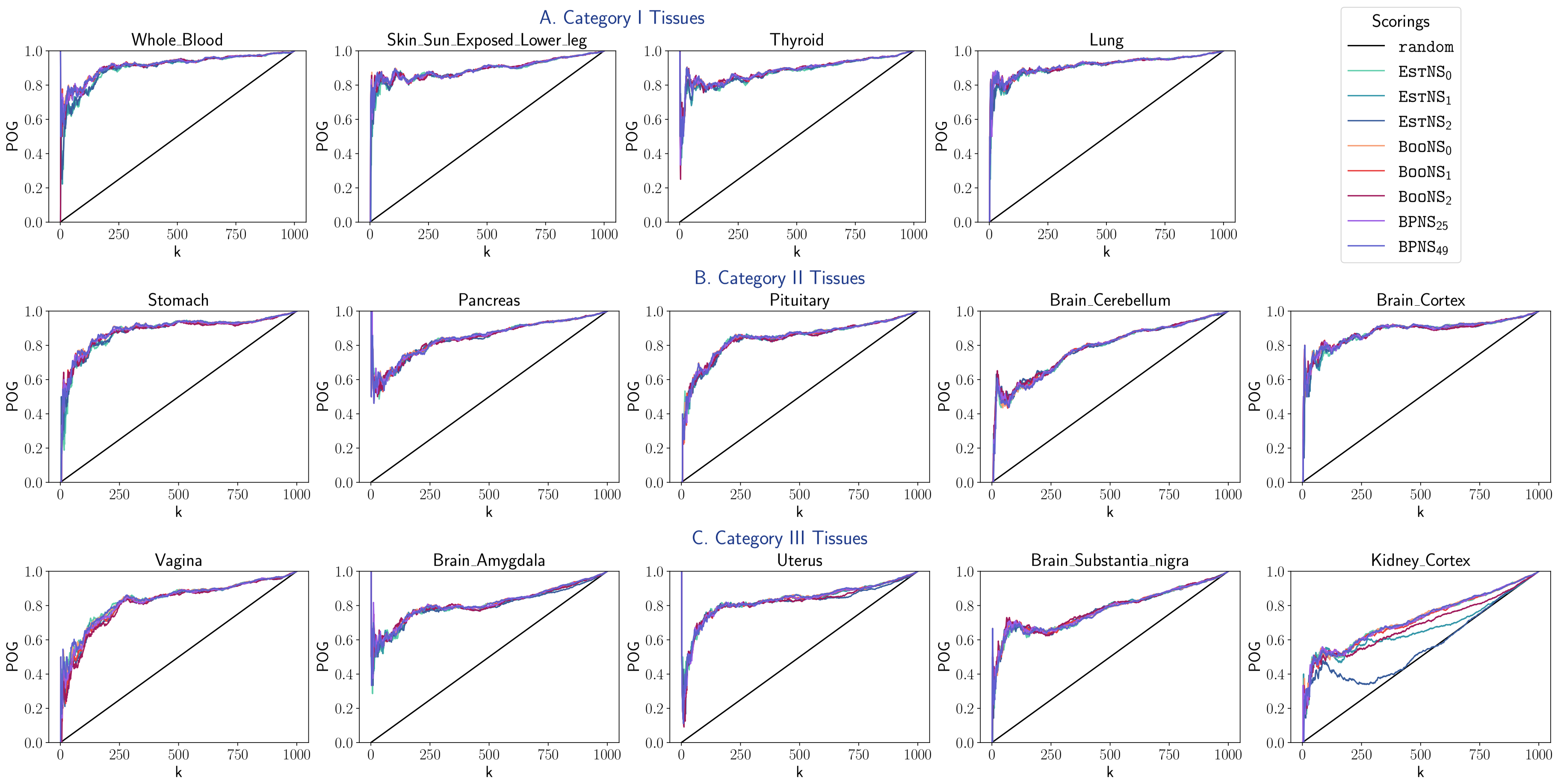


**Suppl. Fig. S7 Replication Analysis in Semi Simulated Dataset of Other Tissues** with replication dataset size = 60% sample size of discovery dataset. Similar CAT plots as in Suppl. Fig. S5 depicting replicability of all scorings.

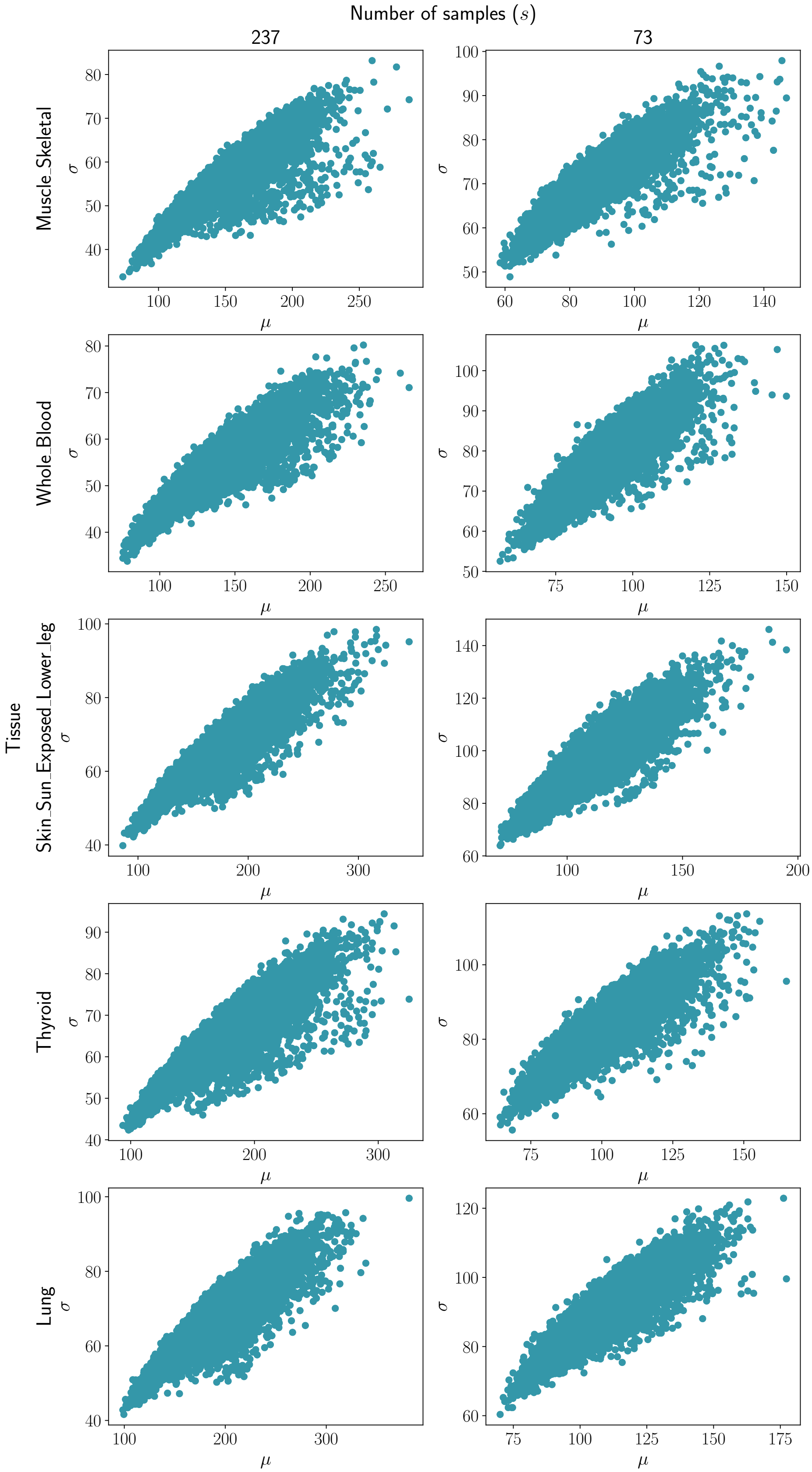


**Suppl. Fig. S8 Validation Analysis using a Real-World Reference Dataset.** Scatter plots depicting the change in $\sigma$ with respect to $\mu$ in degree centrality, and the effect of sample size $s$ on the same across different tissues.

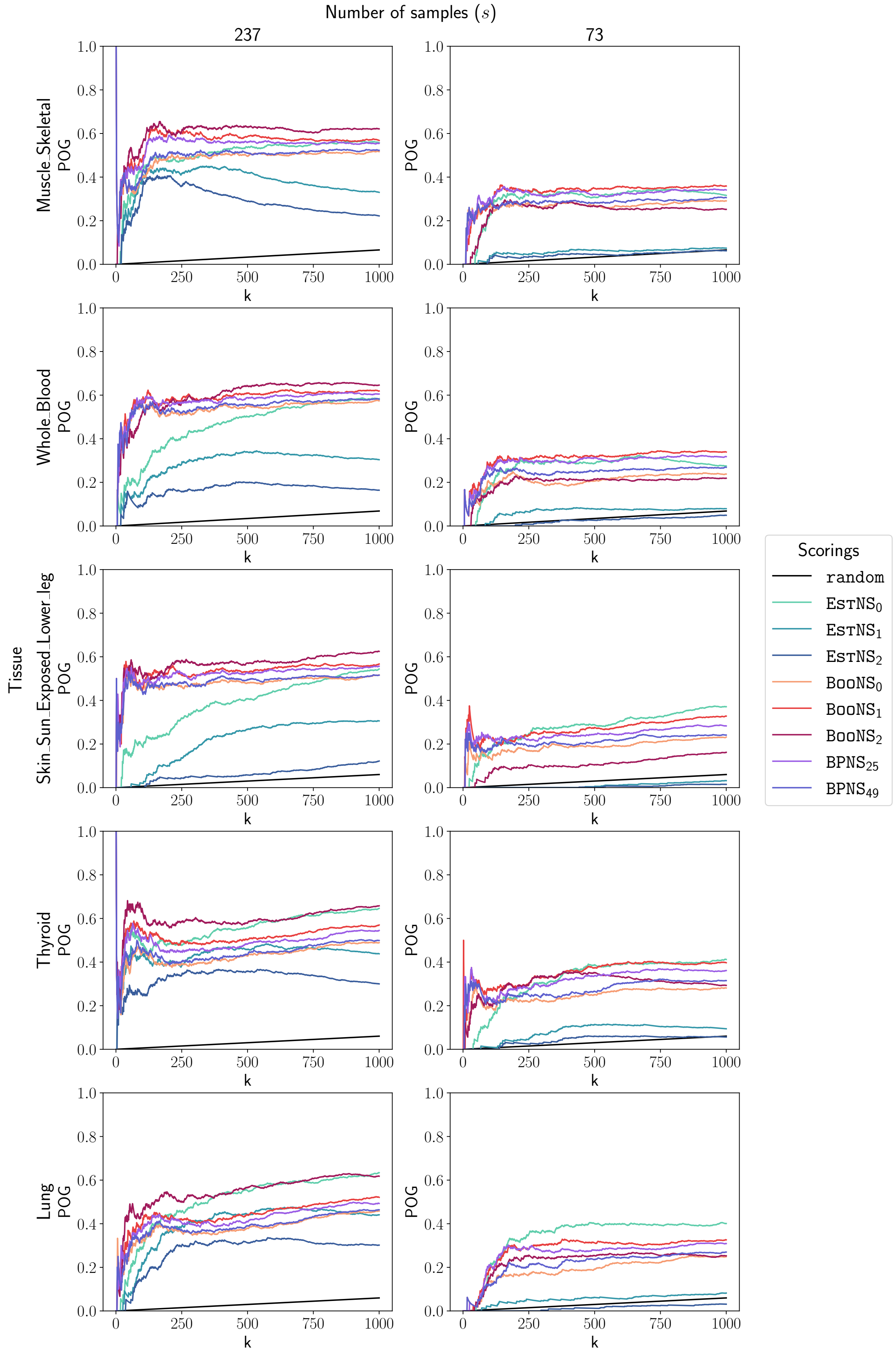


**Suppl. Fig. S9 Validation Analysis using a Real-World Reference Dataset.** CAT plots depicting validation of degree centrality measures by all scorings in subsampled datasets relative to the corresponding actual (preprocessed) dataset for category I tissues.

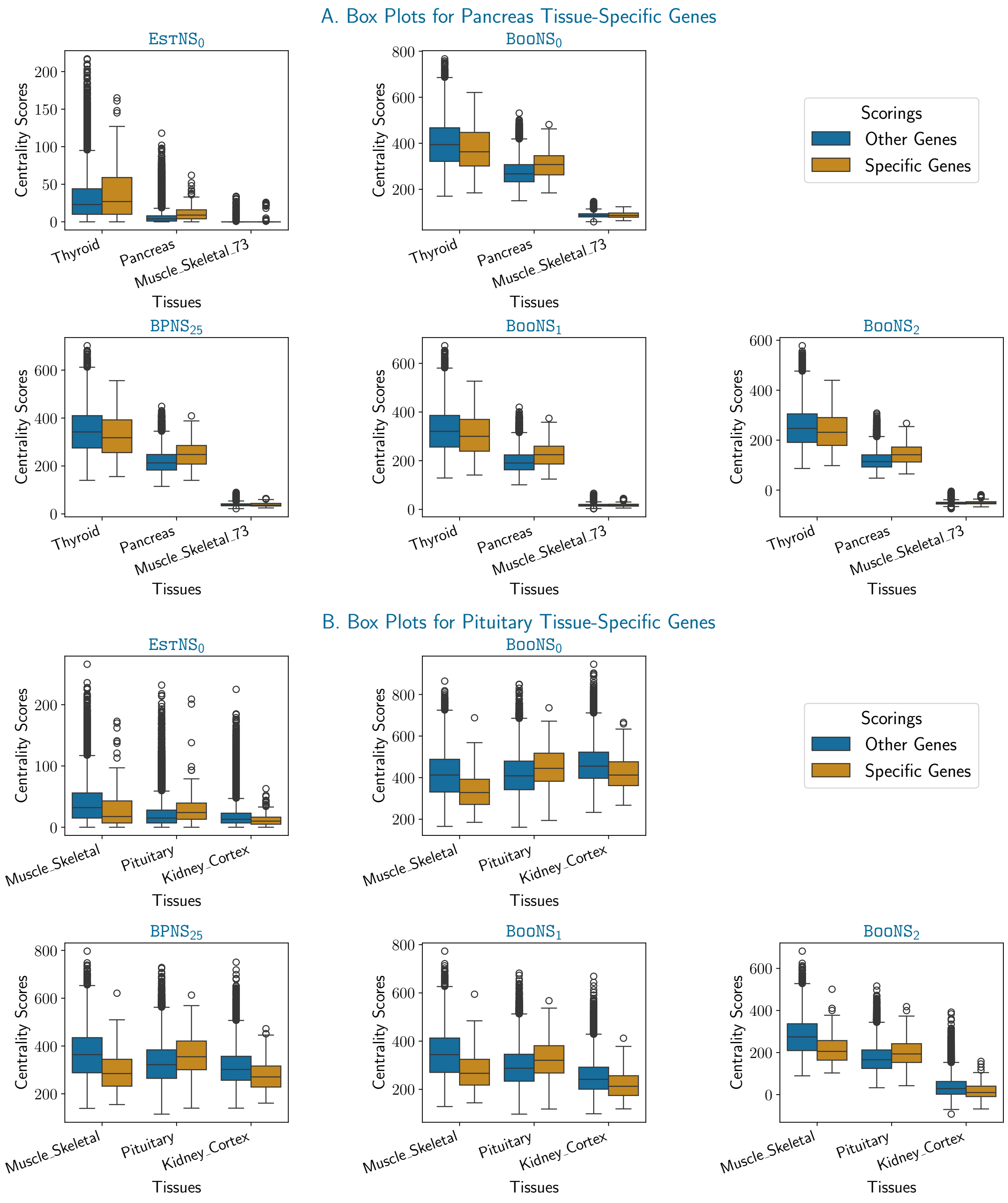


**Suppl. Fig. S10 Tissue Specificity Analysis of Degree Centrality.** A. Box Plots depicting the centrality scores of all the scorings for "Pancreas" tissue-specific genes and all other genes for "Thyroid", "Pancreas" and "Muscle Skeletal (73)" tissues. B. Similar Box Plots depicting the centrality scores of all the scorings for "Pituitary" tissue specific genes and all other genes for "Muscle Skeletal", "Pituitary" and "Kidney Cortex" tissues.

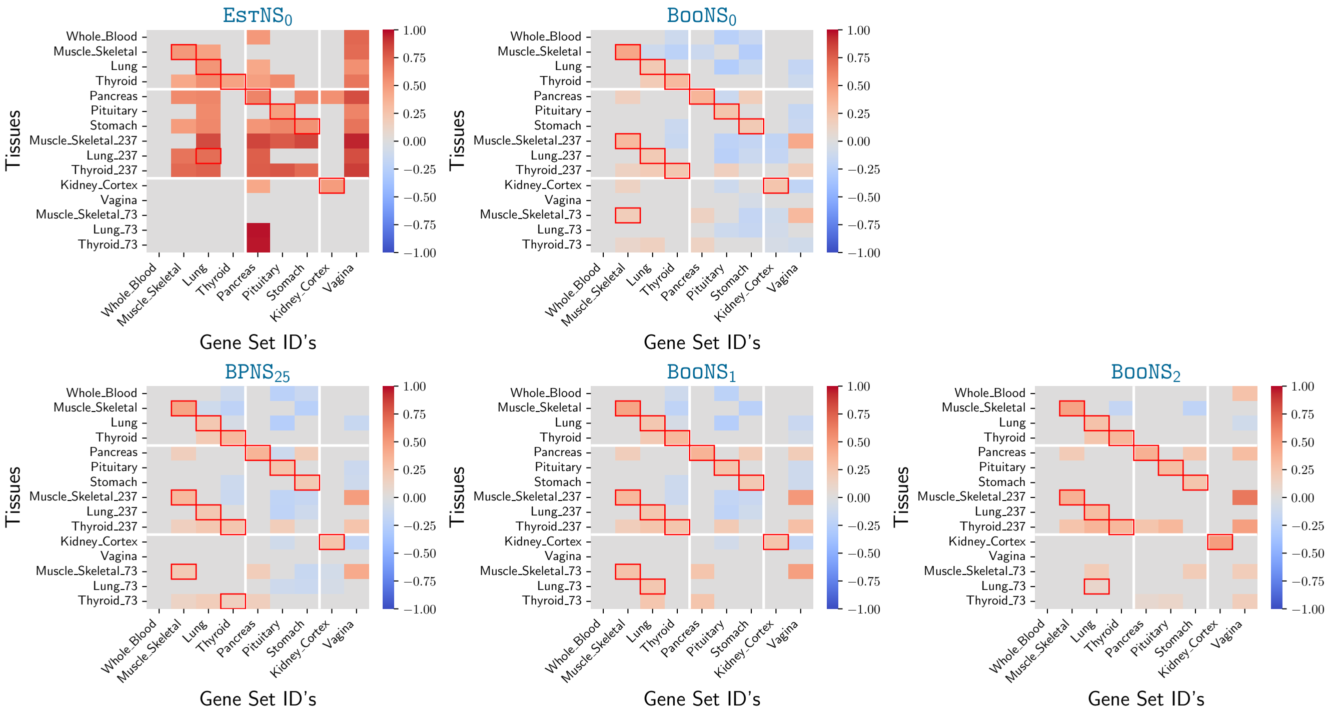


**Suppl. Fig. S11 Tissue Specificity Analysis with 10% FDR threshold of Degree Centrality.** Heatmaps show the enrichment scores of degree centrality measures by all scorings in different tissues, similar to Fig. 6 in main text.

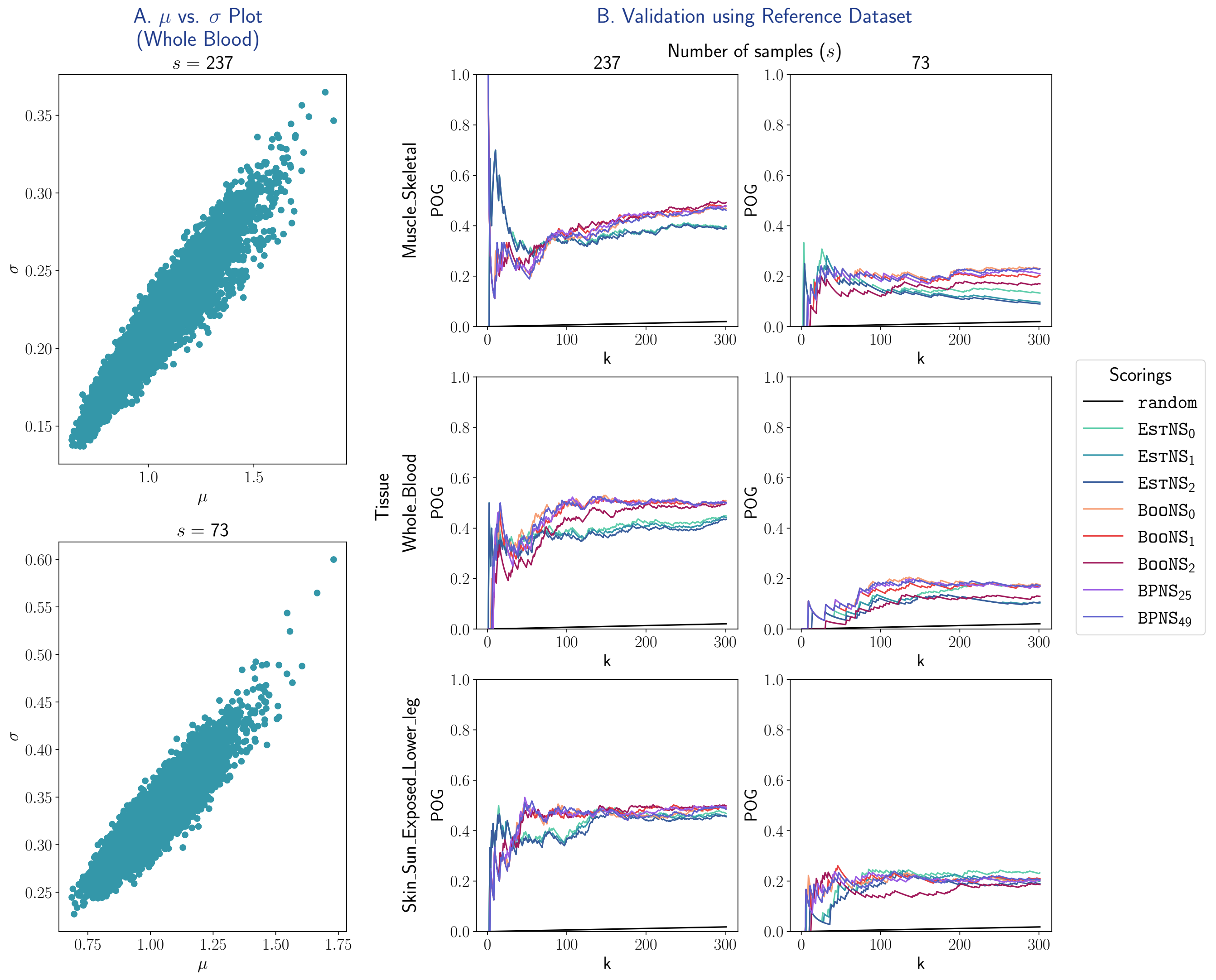


**Suppl. Fig. S12 Validation of PageRank Centrality Scorings using a Real-world Reference Dataset.** A. Plots depicting the change in $\sigma$ with respect to $\mu$, and the effect of sample size $s$ on the same for the Whole Blood tissue. B. CAT plots depicting the validation rate of all scorings in the subsampled datasets relative to the corresponding actual (preprocessed) dataset for Muscle-Skeletal, Whole Blood and Skin tissues.

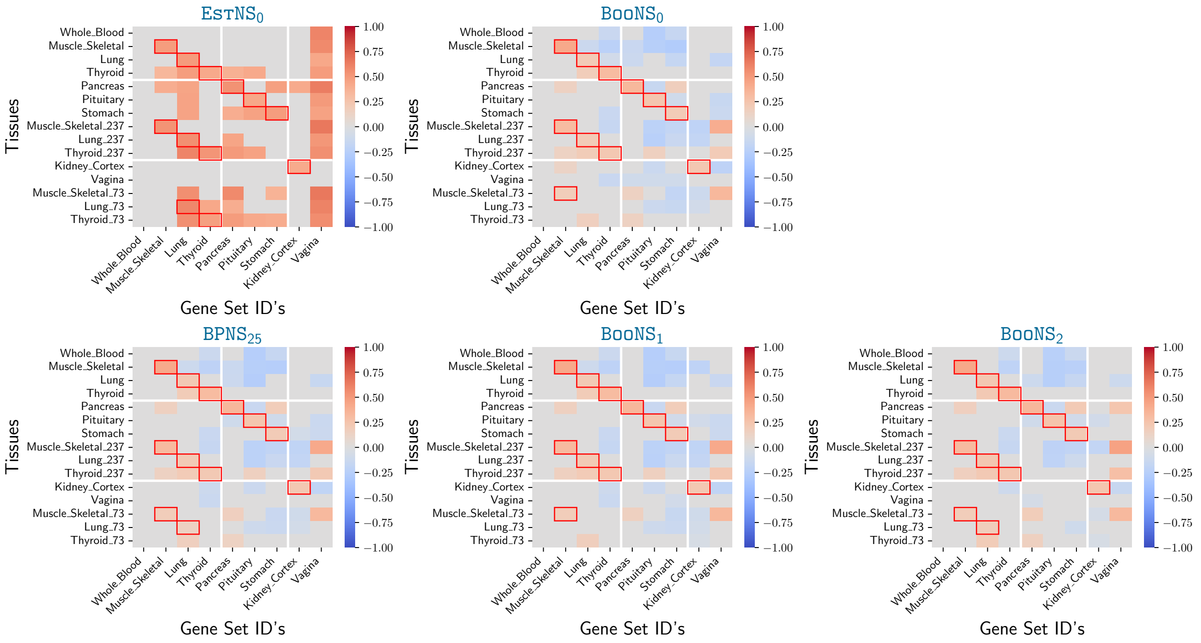


**Suppl. Fig. S13 Tissue Specificity Analysis of PageRank Centrality.** Similar heatmaps, as in Fig. 6 in main text, showing WebGestalt-based enrichment scores of tissues using a $5\%$ FDR threshold.

### C. Supplementary Data/Files

*Suppl. Data S1:*
The original and preprocessed datasets used in the analysis. Link: https://drive.google.com/drive/folders/1pNKQfgAczMcJUzQsTmOgionKpb2LG4zj?usp=sharing

*Suppl. Data S2:*
Scores and rankings in real world datasets (including for $\hat{\theta}$). Link: https://drive.google.com/drive/folders/1XjIoEo4r34P2Yo3mf3HOjvGDmMq_sSvt?usp=sharing

*Suppl. Data S3:*
Hyperparameter tuning – Kendall's $\tau$ values. Link: https://drive.google.com/drive/folders/1sYYxnq3iqmUrGeu6R7UAXCUjeVeQq5P6?usp=drive_link

*Suppl. Data S4:*
POG @ $k$ values for all scoring systems across different scenarios. Link: https://drive.google.com/drive/folders/1GhVQGV8Z1QPM2FCH_Oj7_djX1lukbgwv?usp=sharing

*Suppl. Data S5:*
Recall @ $k$ results across all settings. Link: https://drive.google.com/drive/folders/15FTQ952Cm8VYWHh3_uWo7RN1MiMFL-5a?usp=sharing

*Suppl. Data S6:*
Discovery and replication datasets used for replication analysis on real-world GTEx data. Link: https://drive.google.com/drive/folders/1m-Su6_7VipAPzIu-9WMFLAoa-QlkQrr-?usp=sharing

*Suppl. Data S7:*
Results from the GSEA analysis of $\texttt{BooNS}_1$ based scoring/ranking. Link: https://drive.google.com/drive/folders/1pGZcL6v-BVCa1Vwcs85j-DG4S1g7Nv7P?usp=sharing

*Suppl. Data S8:*
CAT plots of the different analyses performed for PageRank centrality. Link: https://drive.google.com/open?id=1mxa6v799BNAQZl6HHSRyGCz5HJDNglg9&usp=drive_fs
